# The stiffness-sensitive transcriptome of human tendon stromal cells

**DOI:** 10.1101/2021.05.27.445865

**Authors:** Amro A. Hussien, Barbara Niederöst, Maja Bollhalder, Nils Goedecke, Jess G. Snedeker

**Affiliations:** Institute for Biomechanics, ETH Zurich, Zurich, 8092, Zurich, Switzerland; Balgrist University Hospital, University of Zurich, Zurich, 8008, Zurich, Switzerland

## Abstract

Matrix stiffness and its effects on tensional homeostasis act as major regulators of cellular states in health and disease. Stiffness-sensing studies are typically performed using cells that have acquired “mechanical memory” through prolonged propagation in rigid mechanical environments, e.g. tissue culture plastic (TCP). This may potentially mask the full extent of the stiffness-driven mechanosensing programs. To address this, we developed a biomaterial system composed of two-dimensional mechano-variant silicone substrates that is permissive to large-scale cell culture expansion processes. We broadly mapped the stiffness-mediated mechano-responses by performing RNA sequencing on human tendon-derived stromal cells. We find that matrix rigidities approximating tendon microscale stiffness range (*E.* ~35 kPa) distinctly favor programs related to chromatin remodeling and Hippo signaling; whereas more compliant stiffnesses (*E.* 2 kPa) were enriched in responses related to pluripotency, synapse assembly and angiogenesis. We also find that tendon stromal cells undergo dramatic phenotypic drift on conventional TCP, with near-complete suppression of tendon-related genes and emergence of expression signatures skewed towards fibro-inflammatory and metabolic activation. Strikingly, mechano-variant substrates abrogate fibroblasts activation, with tenogenic stiffnesses inducing a transcriptional program that strongly correlate with established tendon tissue-specific signatures. Computational inference predicted that AKT1 and ERK1/2 are major signaling hubs mediating stiffness-sensing in tendon cells. Together, our findings highlight how the underlying biophysical cues may dictate the transcriptional identity of resident cells, and how matrix mechano-reciprocity regulates diverse sets of previously underappreciated mechanosensitive processes in tendon stromal fibroblasts.

## 1. Introduction

Dysregulated mechanics is a hallmark of tendon pathology. Aberrant extracellular matrix stiffening is a feature of progressive aging, and is associated with several chronic inflammatory and metabolic disorders. Stiffened microenvironments in which tendon stromal cells reside significantly differ from that of healthy tissues, with growing body of evidence suggesting that matrix stiffness cues govern key cellular processes.^[1,2]^ Extracellular matrix stiffening is a driver of persistent activation of stromal fibroblasts. Resident stromal cells secret matrix constituents during tissue development and sustain it during homeostasis. In return, extracellular matrix niches relay active and passive biophysical cues that are transduced intracellularly into biochemical signals via mechanotransduction machineries.^[3,4]^ Enhanced matrix stiffness acts as a feed-forward activation loop which enhances fibro-inflammatory activation of stromal cells leading to excessive tissue remodeling and further stiffening of the extracellular matrix.^[5,6]^ How resident fibroblasts sense and integrate matrix stiffness cues is a fertile area for investigation.

Standard tissue culture polystyrene (TCP) vessels have been routinely used for adherent cell culture for over five decades.^[7]^ Although TCP surfaces are optically clear and facilitate cell adhesion *in vitro*, they have supra-physiological levels of stiffness with elastic modulus in the range of Gigapascals (GPa). In contrast, most connective tissues have cell-scale mechanical stiffness with *E.* modulus values in the kilopascal (kPa) range, which may increase by up to two orders of magnitude in the case of fibrosis.^[1,8]^ It is now widely acknowledged that progressive passaging on TCP mechanically activate stromal fibroblasts *in vitro* and instill epigenetically-imprinted mechanical memory that is retained over long-term. Numerous biomaterial systems have been developed to closely mimic optimal tissue stiffness.^[9]^ Previous mechanobiology studies have largely harnessed polyacrylamide hydrogels or Sylgard^®^-based silicone substrata in small to medium scales limited to experimental work. Although polyacrylamide hydrogels have been the most widely employed substrates in rigidity-sensing investigations, these require complex, multistep protocols to facilitate hydrogels bonding to glass slides or to conjugate ECM proteins to its surfaces. Furthermore, increasing the polyacrylamide stiffnesses by varying the crosslinking density simultaneously alter hydrogel pore size and eventually ECM ligand tethering.^[10]^

Alternatively, silicone-based substrates are traditionally fabricated using conventional Polydimethylsiloxane (PDMS) Sylgard^®^ kits. Sylgard 184 formulation was originally designed as a filler to insulate sensitive electrical/electronic circuits, and it contains significant amounts of silica nanoparticle impurities.^[11]^ This significantly contributes to its batch-to-batch variability and reproducibility issues, especially when Sylgard 184 parts are mixed in non-stoichiometric ratios (e.g. 80:1 or 100:1) to produce cultures substrates at the softer end of the spectrum (~ 1-2 kPa).^[12]^ There is clearly a need for an *in vitro* system that enables rigidity-sensing mechanotransduction studies, while seamlessly integrates within standard cell culture practices to facilitates large-scale cell expansion in tissue-like physiological stiffnesses. Here, we introduce a mechano-variant silicone platform with tunable stiffness spanning a wide range of connective tissue-like moduli (2 - 180 kPa). We used this system to map the stiffness-induced transcriptional mechano-responses of tendon-derived stromal cells.

## 2. Results

### 2.1 Tunable mechano-variant PDMS substrates for mechanobiology

To overcome the limitations associated with using conventional mechano-culture substrates, we developed a tunable PDMS mechano-variant culture system using pure, room temperature vulcanizing (RTV) silicone. We started from vinyl-terminated (PDMS), hydride-terminated PDMS, and MethylHydrosiloxane copolymer cross-linker which react in the presence of a platinum catalyst **(Figure 1A)**. We stoichiometrically balanced the silicone precursors to yield two separate components (Part A and Part B) that start to polymerize when mixed at a 1:1 mixing ratio. This design criterion is especially critical for the consistent reproducibility of the stiffness of soft substrates (< 5kPa), which is a bottleneck when commercial Sylgard^®^ 184 kits are used. Since the silicone mixture precursors start reacting immediately when Part A and Part get in contact, we further optimized the time between mixing and placing the mixture on the hotplate to tune the target stiffness of the bulk material **(Figure 1B)**. Additionally, we co-polymerized a fixed amount of undecenoic acid to grafted carboxyl groups in the PDMS surfaces, thus providing tethering points for covalent conjugation of the primary amines of ECM proteins.

**Figure 1.**
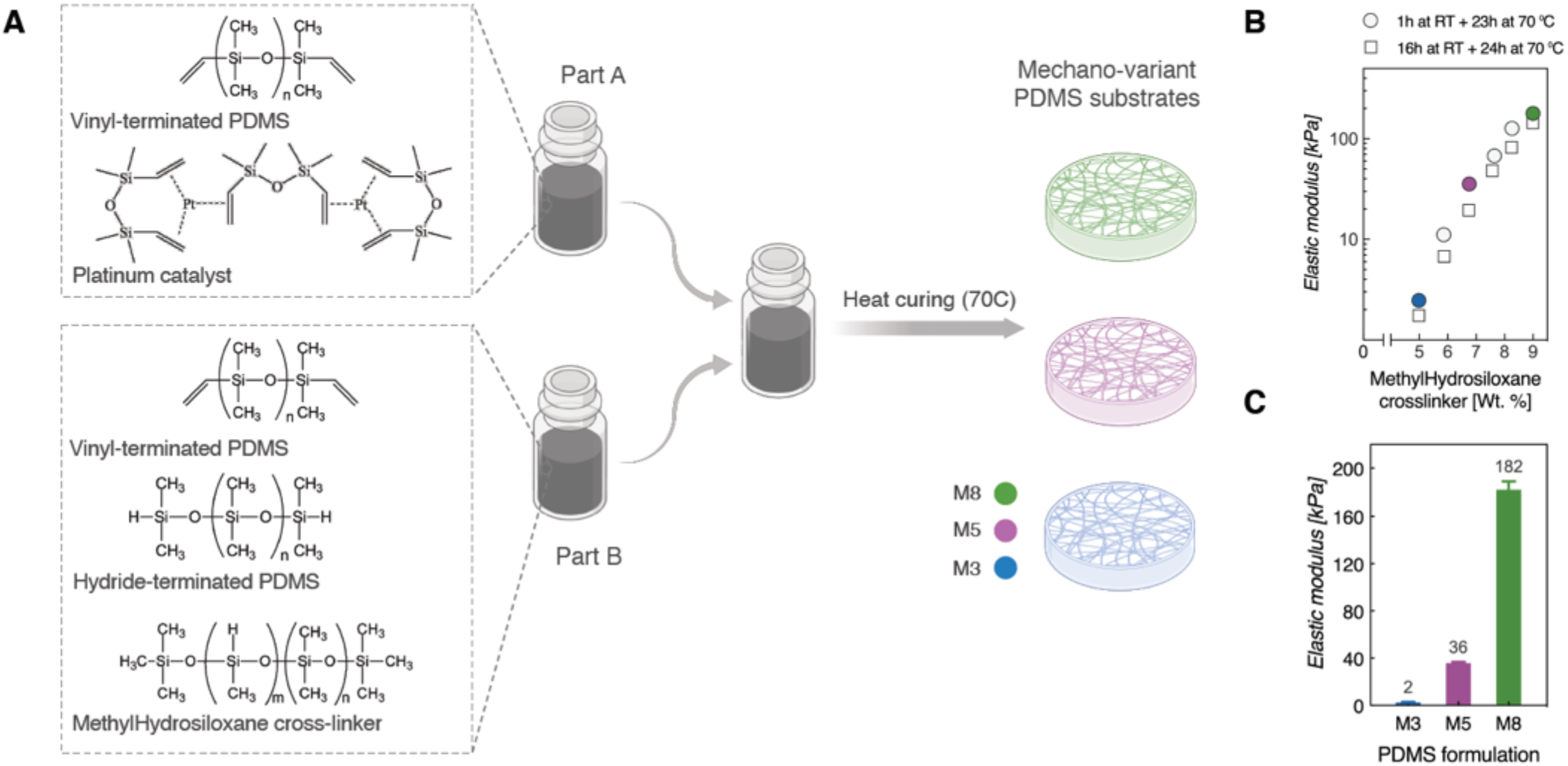
Tunable mechano-variant PDMS substrates for mechanobiology. The silicone formulations cover wide-range of elasticities from very soft to stiff ranges. **(A)** Schematic illustration of the constituent components of the PDMS culture substrates. Recipes were formulated so that the optimal polymerization conditions are initiated by mixing Part A and Part B at 1:1 mixing ratio, while tuning the target *E.* modulus only by varying the amounts of MethylHydrosiloxane copolymer cross-linker in Part B. **(B)** Material characterization of substrates mechanical properties (*E.* modulus) for the indicated PDMS formulations, as measured by micro-indentation using a calibrated piezoresistive Femto-Tools™ probe. PDMS stiffness is dependent on the MethylHydrosiloxane cross-linker concentration and curing conditions. **(C)** PDMS stiffness ranges used in this study. Bars represent the mean ± SD of the closed symbols in figure (B).

This step permits the use of EDC-NHS amine-reactive crosslinker to couple collagen I to the PDMS substrates. Ultimately, we successfully produced mechano-variant silicone substrates with elastic moduli (*E*) ranging from 2kPa to 180kPa, spanning the tenogenic stiffness range **(Figure 1C)**.^[13,14]^

### 2.2 Transcriptomic comparison of stiffness-sensing in human tendon-derived stromal cells

To address the role of matrix stiffness in shaping early transcriptional responses of “naïve” tendon-derived stromal cells, we seeded acutely isolated cells directly on collagen type I-coated PDMS substrates at three different stiffnesses: soft (2 kPa), intermediate (35 kPa) and rigid (180 kPa) **(Figure 2A)**. We have previously shown that intermediate stiffnesses (20-40 kPa) favored tenogenic differentiation of bone marrow mesenchymal stromal cells in an ECM ligand-dependent manner.^[13,15]^ Here, tendon-derived stromal cells were not expanded on TCP and were maintained in culture until reaching 70-80% confluency. We first conducted pairwise differential expression analysis which revealed progressive increase in the number of differentially expressed genes (DEGs) at the extreme of stiffnesses (2kPa *vs.* 180kPa: 259 DEGs) compared to the comparisons within narrower ranges (2kPa *vs.* 35 kPa: 40 DEGs or 35kPa *vs.* 180kPa: 100 DEGs) **(Figure 2B and C)**. DEGs showed minimal overlapping across the stiffness comparisons suggesting that stiffness-mediated transcriptional responses might be unique across each stiffness range **(Figure 2C)**.

**Figure 2.**
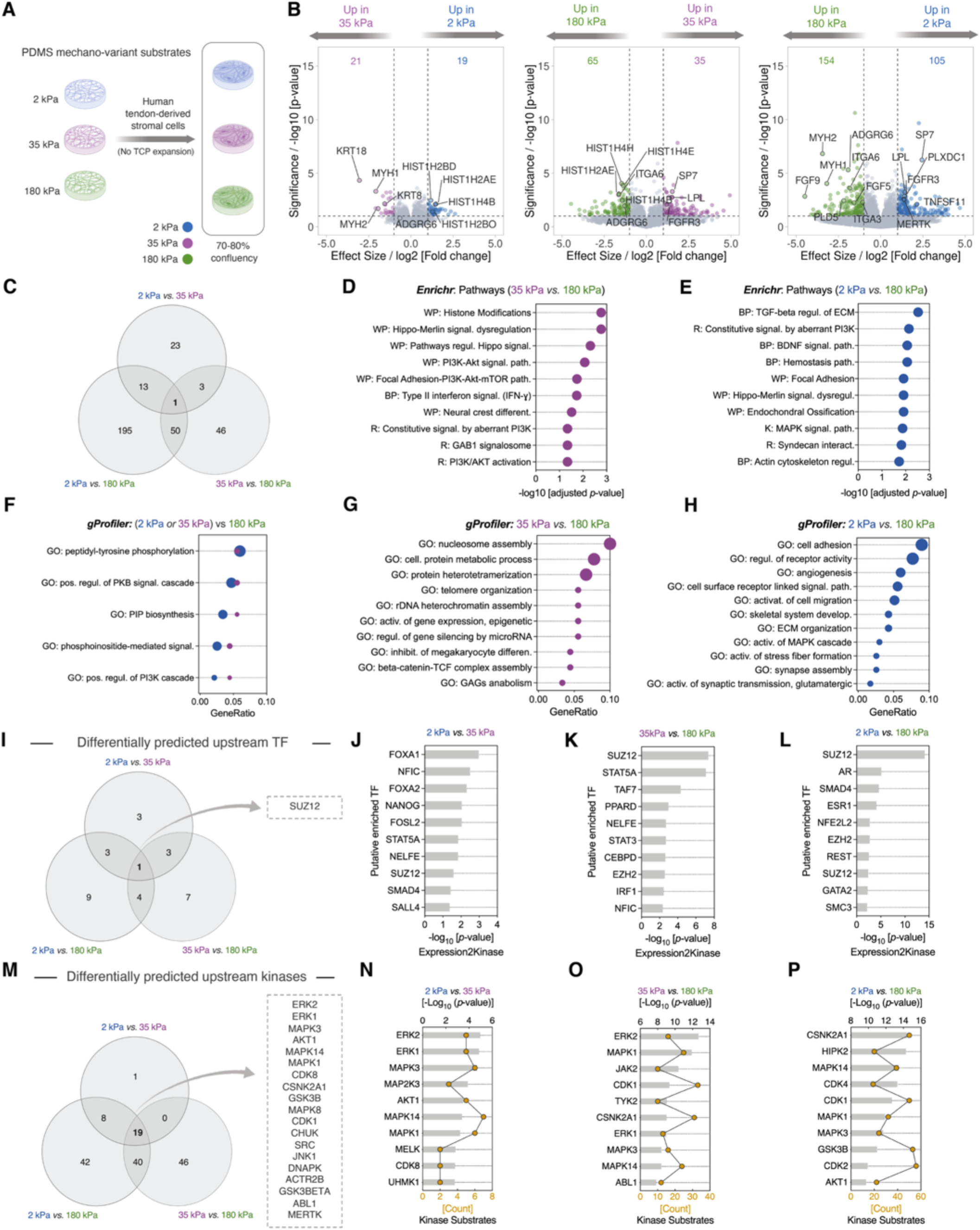
Transcriptomic comparison of stiffness-sensing in human tendon-derived stromal cells. **(A)** Schematic overview of the range of substrate stiffnesses that were used to interrogate stiffness-sensing in this study. Healthy human tendon-derived cells were isolated from patients who underwent autograft tendon transfer for the repair of knee anterior cruciate ligament (ACL). Dissociated cells were directly seeded on PDMS substrates without the standard expansion on tissue culture plastics. **(B)** RNA-seq volcano plots of DEGs across different stiffness comparisons. Differential expression was determined by *DESeq2* method. Colored values represent significant DEGs, which are defined with cut-off values of |fold change| ± 1 and *p*-value < 0.01. **(C)** Venn diagram depicting the overlapped significantly expressed genes (DEGs) between different stiffness comparisons. **(D** and **E)** Top 10 enriched pathways for each stiffness comparison; (D) 35kPa *vs.* 180kPa, (E) 2kPa *vs.* 180kPa. Enrichment analysis was performed in *Enrichr* using both up-regulated and down-regulated genes. DEGs was interrogated against BioPlanet 2019 (BP), Reactome (R), WikiPathways 2019 (WP), KEGG 2019 (K) databases. The list of significantly enriched pathways was manually curated, and it is listed according to the smallest *p*-values obtained across all the databases. **(F)** Shared enriched biological processes (BP) GO terms between the indicated stiffness comparisons using *gProfiler* over-representation analysis of both upregulated and downregulated DEGs. **(G** and **H)** Top enriched biological processes (BP) GO terms in each stiffness comparison; (G) 35kPa *vs.* 180kPa, (H) 2kPa *vs.* 180kPa. Over-representation analysis was performed using a hypergeometric over-representation test against the GO database with both upregulated and downregulated DEGs. Significance achieved at *q-*value < 0.05. **(I)** Venn diagram showing the overlapped predicted upstream transcription factors (TF) between the different stiffness comparisons. **(J-L)** Bar plots show the predicted top 10 most significantly enriched transcription factors upstream of the DEGs. Predicted TFs are sorted by significance level (adjusted *p*-value < 0.05). **(M)** Venn diagram depicting the overlapped predicted upstream kinases between the different stiffness comparisons. **(N-P)** Bar plots indicate the predicted top 10 most significantly enriched kinases upstream of the DEGs. Predicted kinases are ranked by significance level (adjusted *p*-value < 0.05). TF and kinases were predicted using the TF and Kinase Enrichment Analysis modules of the *Expression2Kinase* tool.

Using *Enrichr*, we next searched for stiffness-responsive pathways that are functionally enriched in the up-regulated and down-regulated DEGs.^[16–18]^ We found that the intermediate stiffness range (35 kPa) *vs.* the stiffer substrates (180 kPa) was enriched in pathways known to be mechanosensitive or mechanically regulated, such as chromatin and histone modifications, Hippo-Merlin pathway, and the PI3K-Akt pathway **(Figure 2D)**. In contrast, stromal cells on softer stiffnesses (2 *vs.* 180kPa) were transcriptionally enriched in a wider-range of pathways related to extracellular matrix (TGF-beta regulation of ECM, syndecan interactions), endochondral ossification, cell adhesion and cytoskeleton **(Figure 2D)**. Of note is that the PI3K-Akt pathway and its related processes were consistently enriched across the two stiffness comparisons, suggesting it might be a key signaling hub regulating the stiffness-sensitive response in tendon stromal fibroblasts **(Figure 2F)**. Examining the enriched pathways in the stiffness-specific genes, *i.e.* DEGs exclusively expressed in one stiffness range but not the others, revealed that histone modifications and TGF-beta regulation of ECM were the most significantly enriched pathways in the intermediate (35 *vs.* 180kPa) and soft stiffnesses (2 *vs.* 180kPa), respectively **(Supplementary figure S1)**. This suggest that the top enriched pathways in each stiffness comparison are largely driven by a subset of stiffness-sensitive transcripts among the top differentially expressed genes.

To further characterize the enriched transcriptional processes, we performed overrepresentation analysis (ORA) using Gene Ontology biological processes (GO-BP) database. Again, biological processes related to histone modifications, chromatic remodeling and epigenetics were significantly enriched in intermediates stiffnesses, with seven out of the top 10 overrepresented GO-BP terms **(Figure 2G)**. Consistent with the pathway analysis, wider range of GO terms were overrepresented in soft substrates with cell adhesion and migration, angiogenesis and synapse assembly terms among others occupying the top 10 most enriched terms **(Figure 2H)**.

Next, we sought to computationally deduce the upstream regulators that are likely responsible for the observed transcriptional programs or display an altered activity in response to the underlying substrate stiffness. We used a complimentary bioinformatic approach based on the *Expression2Kinase* (X2K) algorithm, which rely on the global pattern of transcriptome changes to construct a protein-protein interaction network in order to infer potential upstream regulators.^[19]^ We performed a transcription factor enrichment analysis on each of the three pairwise comparisons **(Figure 2C and I)**. In the soft-to-intermediate stiffness comparison (2 *vs.* 35 kPa), we noted significant enrichment of transcriptional regulators related to endodermal/mesodermal differentiation (FOXA1, FOXA2, and SMAD4) and self-renewal and pluripotency (NANOG and SALL4) **(Figure 2J)**. In contrast, intermediate-to-hard (35 *vs.* 180kPa) and soft-to-hard (2 *vs.* 180kPa) comparisons were consistently enriched in TFs members of Polycomb repressive complex 2 (SUZ12 and EZH2) which are known to function as heterochromatin modifiers. Intriguingly, intermediate-to-hard (35 *vs.* 180kPa) range uniquely showed enrichment of TFs related inflammation and cytokines signaling (STAT3 and IRF1) and adipogenic differentiation (PPARD and CEBPD) **(Figure 2K and L)**.

In the next step, we extended our analysis and used the DEGs and predicted kinases as seed inputs for the kinase enrichment analysis (KEA) module of the X2K tool **(Figure 2M)**. This analysis showed that mitogen-activated protein (MAPK) kinases (ERK1/MAPK3, ERK2/MAPK1, and p38α/MAPK14) and the serine/threonine-protein kinase (AKT1) consistently ranked among the top 10 most enriched kinases across the different pairwise comparisons **(Figure 2N-P)**. All of which have been implicated in mechano-signaling, in some form or another, in the context of stromal connective tissues.^[20]^ Specifically, we recently demonstrated that ERK1/2 acts a signaling checkpoint mediating the loss of mechanical tension in tendons.^[21]^

Furthermore, X2K algorithm predicted distinct kinases that were enriched in a stiffness-specific manner. While Janus kinase (JAK) family members (JAK1, JAK2, JAK3, and TYK2) were enriched in intermediate-to-hard (35 *vs.* 180kPa) stiffness range, Cyclin-dependent kinases (CDK1, CDK2, CDK4, and CDK6), Casein kinases, and HIPK kinases (HIPK1 and HIPK2) subfamilies were substantially more enriched in soft-to-hard (2 *vs.* 180kPa) stiffnesses comparison **(Figure 3N-P, Supplementary figures S2–4)**. This suggests that matrix stiffness potentially alters tendon stromal cells signaling through convergent (stiffness-sensitive) and distinct (i.e. stiffness-specific) pathways.

**Figure 3.**
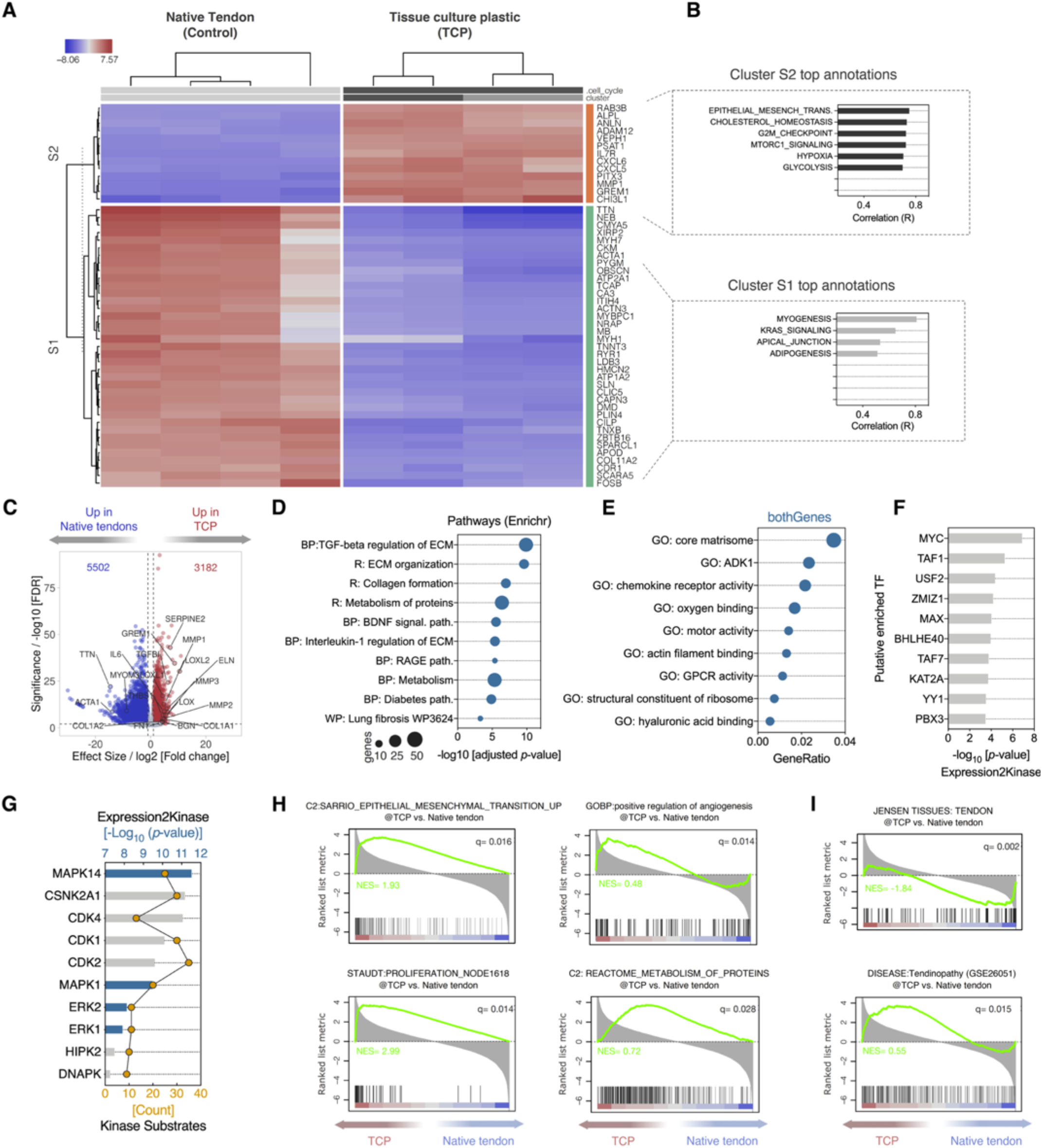
Global transcriptomic characteristics of tenocytes phenotypic drift in standard TCP culture conditions. (**A**) Heatmap of gene-level hierarchical clustering of the top 50 differentially expressed genes (DEG) in tendon stromal fibroblasts cultured in tissue culture plastic (TCP) vs. vehicle control. Columns represent individual samples (N=4 biological replicates from different donors). Blue denotes downregulated genes; red denotes upregulated genes. **(B)** Functional annotation of S1 and S2 gene clusters in the heatmap (encompassing top 500 DEGs). Bar plots depict the fisher-weighted, average correlations of the cluster with the annotation terms queried against MSigDB Hallmark database. **(C)** RNA-seq volcano plot of DEGs of TCP-cultured human tenocytes relative to native tendons. Colored dots show the 8,684 significantly expressed genes, as determined by *DESeq2* methods, with the horizontal line corresponding to an FDR ≤0.01 and vertical lines are at a cutoff of log2[Fold change] ± 1. **(D)** Enriched pathways analysis of a subset of DEGs using *Enrichr* queried against BioPlanet 2019 (BP), Reactome (R), WikiPathways 2019 (WP), and KEGG 2019 (K) Human databases. All hits had an adjusted *p*-value < 0.05. **(E)** Overrepresentation analysis (ORA) of Molecular Functions GO terms of upregulated and downregulated DEGs. Analysis was performed using a hypergeometric over-representation test against the GO database, with significance achieved at *q-*value < 0.05. **(F)** Bar plot depicts the predicted top 10 most significantly enriched transcription factors upstream of the DEGs. Predicted TFs are sorted by significance level (adjusted *p*-value < 0.05). **(G)** Top 10 kinases upstream of predicted TFs in (F) were identified using the Kinase Enrichment Analysis module of the *Expression2Kinase* pipeline. **(H)** GSEA enrichment plots of some of the top differentially expressed gene sets. The plot’s black vertical lines represent the ranked gene hits in the gene set. The green line represents the normalized enrichment score (NES). The more the NES curve is skewed to the upper left of the plot, the more the gene set is enriched in the TCP group. In contrast, when the curve shifts to the lower right, this reflects a more enrichment in the Native Tendon control group. The red-to-blue colored scale at the bottom represents the degree of correlation of genes in the TCP group with the phenotype in the gene set (Red: positive correlation; blue: negative correlation) or *vice versa* for the Native Tendon control group. Note the significant regulation of gene set terms related to EMT, cellular proliferation, metabolism, and angiogenesis. **(I)** GSEA enrichment plots of tendon-specific (Top) and tendinopathy (bottom) signatures.

### 2.3 Global transcriptomic characteristics of tenocytes phenotypic drift in standard TCP culture

Prolonged culture expansion on conventional tissue culture plastic (TCP) activates pro-fibrogenic transcriptional and epigenetic programs that are imprinted as “mechanical memory” in stromal cells.^[22–24]^ In addition, it has been reported that human tendon-derived stromal fibroblasts undergo phenotypic drift when explanted and progressively passaged *in vitro* in the ultra-stiff TCP.^[25]^ To assess the full extent of early transcriptomic responses of the TCP-induced phenotypic drift, we initiated tendon stromal fibroblasts cultures from freshly isolated cells on conventional culture vessels without ever passaging the cells. When cells reached 70-80% confluency (approximately after 5-7 days), we collected the cells for bulk RNA sequencing and compared genome-wide mRNA transcription to native, tissue-level controls. Details on clinical information of all donors are included in Supplementary Information **(Supplementary Table S1)**.

Differential expression analysis revealed strong and significant regulation of 8,684 DEGs between TCP cultures and native tendons among which 5,502 were upregulated and 3,182 were downregulated at a threshold of |fold change (FC)| ± 1 and adjusted *p* < 0.01 **(Figure 3 A-C, Supplementary figures S5)**. Notably, differentially upregulated genes in TCP substrates showed positive correlation with reference database MSigDB Hallmarks gene sets: epithelial-mesenchymal transition (EMT), hypoxia, and metabolism [represented by cholesterol homeostasis, MTORC1 signaling and glycolysis] **(Figure 3B)**. In contrast, Myogenesis gene set showed the strongest correlation (Pearson *r* = 0.81) with upregulated genes in the Native tendon contrast, possibly reflecting the overlap between tendon and muscle transcriptomes.

To systematically characterize the enriched transcriptional programs, we performed pathway enrichment analysis on DEGs of S2 subcluster using the tool EnricrR.^[16,17]^ Approximately half of the top 10 enriched hits were pathways related to fibro-inflammatory regulation of the ECM, while 4/10 of the enriched terms are representative of metabolic pathways **(Figure 3D)**. We then carried out an over-representation analysis (ORA) to examine what Gene Ontology (GO) terms are over-represented in all the up-regulated and down-regulated DEGs.^[26]^ GO terms related core matrisome topped the over-represented gene sets, with consistent enrichment of terms related to chemokine activity, cytoskeletal remodeling, and GPCR signaling **(Figure 3E)**.

Next, we sought to delineate the regulatory signaling kinases and transcriptional factors upstream of the DEGs. X2K TF module identified 32 TFs that may potentially regulate the DEGs **(Figure 3F)**. The top 10 TFs were dominated by transcriptional regulators known to be involved in cell cycle progression, apoptosis, or cellular differentiation (MYC, MAX, BHLHE40) or chromatin remodeling (KAT2A and YY1). Further, X2K kinase enrichment analysis predicted a total of 98 significant kinases (Hypergeometric *p*-value of < 0.05), of which the top 10 most significant kinases are dominated by mitogen-activated protein kinases (MAPK14, ERK1, ERK2) and cyclin-dependent kinases (CDK1, CDK2, CDK4) **(Figure 3G, Supplementary figures S6)**. Finally, to get a global unbiased insight on the potential phenotypes associated with these altered gene programs, we carried out pre-ranked Gene Set Enrichment Analysis (GSEA).^[27,28]^ We found that epithelial-mesenchymal transition (NES 1.93), proliferation (NES 2.99), regulation of angiogenesis (NES 0.48) and metabolism of proteins (NES 0.72) gene sets among the most significant signatures with positive correlation to TCP cultures **(Figure 3H)**. Interestingly, TCP transcriptome showed significant enrichment in tendon-related signatures, with negative correlation with Jensen Tissues (Tendon) and positive correlation with DISEASE: Tendinopathy signatures **(Figure 3I)**.

Collectively, this data provides strong evidence that culture expansion in conventional TCP polystyrene vessels may contribute to fibro-inflammatory and metabolic activations of tendon-derived fibroblasts, with the divergence of the whole transcriptome towards a pathological signature closely resembling that of tendinopathic tendons.

### 2.4 Tenogenic mechano-variant substrates abrogate TCP-mediated inflammatory programs and favor tendon-specific expression signature

Next, we sought to assess how mechano-variant silicone substrates modulate early transcriptional responses of tendon stromal cells in comparison to ultra-stiff TCP. We compared transcriptome profiles of soft, intermediate, and rigid-cultured tendon cells to conventional TCP surfaces **(Figure 4, Supplementary figures S7 and S8)**. Here, we will only highlight the results from the intermediate (35 kPa) stiffness in the interest of maintaining the focus of the paper on tendon-related responses. When cells were conditioned to tenogenic PDMS substrates (*E*. 35 kPa), unsupervised genes subclusters showed consistent and negative correlations to Hallmark annotations related to inflammation (e.g. TNFA Signaling via NFKB, IL6 JAK STAT3, Inflammatory Response) and Myogenesis in the MSigDB Hallmark database **(Figure 4A and B)**. Differential analysis revealed 2,787 significantly expressed genes with 1,112 upregulated and 1,675 downregulated genes at a threshold of |fold change (FC)| ± 1 and adjusted *p* < 0.01 **(Figure 4C)**. Using *Enrichr*, pathway analysis uncovered domination of fibro-inflammatory related pathways such as TGF-β1 regulation of ECM, collagen formation, IL-4 regulation of apoptosis and ECM organization **(Figure 4D)**. Genes overlapping with PI3K-Akt signaling pathway were also significantly enriched, in consistence with the comparison across the PDMS stiffnesses **(Figure 2F)**.

**Figure 4.**
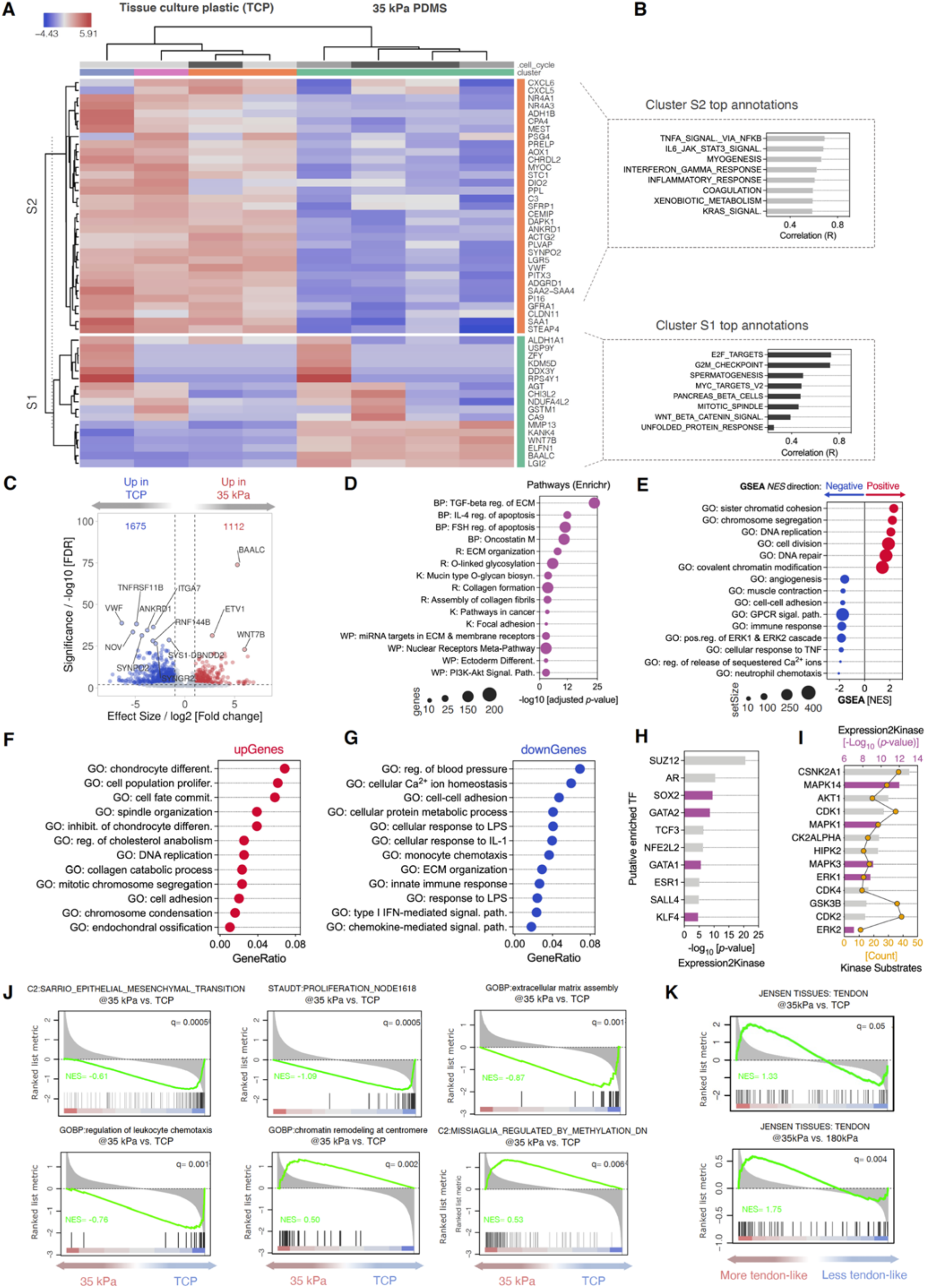
Tenogenic mechano-variant substrates abrogate TCP-mediated inflammatory programs and favor tendon-specific expression signature. (**A**) Heatmap of gene-level hierarchical clustering of the top 50 differentially expressed genes (DEG) in tendon stromal fibroblasts cultured in tenogenic substrates (*E.* ~35kPa) vs. TCP control. Columns represent individual samples (N=4 biological replicates from different donors). Blue denotes downregulated genes; red denotes upregulated genes. **(B)** Functional annotation of S1 and S2 gene clusters in the heatmap (encompassing top 500 DEGs). Bar plots depict the fisher-weighted, average correlations of the cluster with the annotation terms queried against MSigDB Hallmark database. **(C)** RNA-seq volcano plot of DEGs of 35kPa-cultured human tenocytes relative to TCP controls. Colored dots show the 2,787 significantly expressed genes, as determined by *DESeq2* method, with the horizontal line corresponding to an FDR ≤0.01 and vertical lines are at a cutoff of log2[Fold change] ± 1. **(D)** Enriched pathways analysis of a subset of DEGs using *Enrichr* queried against BioPlanet 2019 (BP), Reactome (R), WikiPathways 2019 (WP), and KEGG 2019 (K) Human databases. All hits had an adjusted *p*-value < 0.05. **(E)** Pre-ranked Gene Set Enrichment Analysis (GSEA) of positively and negatively enriched cellular components by Normalized Enrichment Score (NES) in 35kPa cultured cells (Adjusted *p*-value < 0.05). **(F-G)** Overrepresentation analysis (ORA) of Molecular Functions GO terms of upregulated (F) and downregulated (G) DEGs. Analysis was performed using a hypergeometric over-representation test against the GO database, with significance achieved at *q-*value < 0.05. **(H)** Bar plot depicts the predicted top 10 most significantly enriched transcription factors upstream of the DEGs. Predicted TFs are sorted by significance level (adjusted *p*-value < 0.05). **(I)** Top 10 kinases upstream of predicted TFs in (F) were identified using the Kinase Enrichment Analysis module of the *Expression2Kinase* pipeline. **(J)** GSEA enrichment plots of some of the top differentially expressed gene sets. The Plot’s black vertical lines represent the ranked gene hits in the gene set. The green line represents the normalized enrichment score (NES). The more the NES curve is skewed to the upper left of the plot, the more the gene set is enriched in the 35kPa group. In contrast, when the curve shifts to the lower right, this reflects a more enrichment in the TCP control group. The red-to-blue colored scale at the bottom represents the degree of correlation of genes in the 35kPa group with the phenotype in the gene set (Red: positive correlation; blue: negative correlation) or *vice versa* for the TCP control group. Note the significant negative regulation of gene set terms related to EMT, cellular proliferation, ECM assembly, and inflammation. **(K)** GSEA enrichment plots of tendon-specific (Top) and tendinopathy (bottom) signatures.

Preranked GSEA by *fgsea* showed analogous enrichment of GO terms with positive regulation of chromatin remodeling and DNA replication and repair processed **(Figure 4E)**. Interestingly, the enrichment of gene sets related to angiogenesis, ERK1/2 cascades and inflammatory pathways in silicone substrates showed consistent negative gene regulation across all PDMS stiffnesses in comparison to TCP **(Figure 4E, Supplementary figures S7E and S8 E)**. This hints at the possibility that PDMS substrates tend to abrogate the strong TCP-mediated pro-inflammatory and stress responses. Furthermore, ORA analysis showed significant over-representation of GO terms associated with cell fate commitment and musculoskeletal cells differentiation [represented by chondrocyte differentiation, inhibit. of chondrocyte differentiation and endochondral ossification] in the upregulated genes **(Figure 4F)**. In contrast, more diverse GO processes were over-represented in the down-regulated genes dominated by inflammation (seven GOs), in addition to protein metabolism, ECM organization and calcium ion homeostasis **(Figure 4G)**.

Reconstruction of upstream TFs and regulatory kinases deduced consistent enrichment of TFs known to be involved in self-renewal and pluripotency [SOX2, GATA2, GATA1, KLF4] across all PDMS variations relative to TCP, implying that mechano-variant substrates may condition tendon cells towards a more progenitor/stemness-like state. **(Figure 4H, Supplementary figures S7 H and S8 H)**. In addition, SUZ12 was the top deduced TF across all the PDMS comparisons, while ERK1/2 and AKT1 kinases over-populated the list of predicted kinases **(Figure 4H and I)**. Surprisingly, 35 kPa tenogenic substrates induced expression pattern that positively correlated with signatures suggestive of epigenetic processes (chromatic remodeling: NES 0.05, Methylation regulation: NES 0.53) **(Figure 4J)**, while strongly correlated with tendon tissue-specific transcriptomic signature (NES 1.33) **(Figure 4K)**.

## 3. Discussion

In this work, we have established a user-friendly, two-dimensional mechano-variant cell culture system using silicone elastomer gels which is amenable to large-scale cell expansion practices. We have used this new biomaterial system to profile early transcriptional mechano-responses of human tendon-derived stromal cells and defined a stiffness range that reinstate a tendon-like transcriptomic signature in tendon stromal cells *in vitro*. A major motivation behind this study lays in overcoming the long-standing technical challenges in using tunable mechano-variant substrates in routine cell culture practice. We addressed this issue by using commercially available, pure silicones that are devoid of silica nanoparticle fillers.^[29]^ We formulated the constituent precursors to yield balanced two-part liquid components that can be mixed at a 1:1 ratio, irrespective of the target stiffness. With an important step that dramatically simplifies later surface preparation, we co-polymerized undecenoic acid with the reaction mix to introduce carboxylic acids groups that enabled direct covalent coupling of primary amines using the EDC/NHS crosslinking chemistry. Thus, we circumvented the challenge of using the unstable, UV light-dependent sulfo-SANPAH heterobifunctional crosslinker. This allowed us to produce large scale mechano-variant substrates inside sealed tissue culture flasks under sterile conditions. More importantly, we were able to use the platform to propagate freshly isolated tendon fibroblasts at all stiffness ranges (from 2-180 kPa) without exposing the cells to the potentially confounding effects of TCP-mediated mechanical dosing.^[23,30]^

Perhaps one of the most striking findings of this study is that the medium range of stiffness (~35 kPa) conditioned tendon-derived stromal cells transcriptome towards tendon-specific tissue signature. Substrate stiffness-driven tenogenic differentiation of bone marrow stromal cells has been described for many years. Our previous work has established for the first time that intermediate matrix stiffnesses (30-90 kPa) and collagen type I ECM ligands direct differentiation of bone marrow mesenchymal stromal cells towards tendon lineage specification.^[13,15]^ Similarly, Rehmann and colleagues showed that synthetic poly(ethylene glycol)-based elastic substrates promoted expression of tendon lineage-associated genes in human mesenchymal stem cells, with 50kPa modulus being the optimal stiffness.^[14]^

Our results here extend these observations to tendon-derived stromal fibroblasts, and suggest that resident fibroblasts sense the optimal tendon moduli within a narrow range of values approximating that of *in vivo* cell-scale microenvironmental stiffness. Tendon tissue micro- and nanoscale indentation modulus has been characterized, and indeed it has been reported that cell-scale elastic modulus spans a range of tens of kPa, depending on the methods used as well as the anatomical source of the tendon.^[31]^ These findings are consistent with the current view from other biological systems that the optimal mechanical microenvironment for a given cellular behavior *in vitro* often correlates with the corresponding elasticity of the living tissue *in vivo*.^[32]^ In muscles, optimum substrate stiffness that facilitates myotube differentiation and striation matches elasticity of healthy muscle at *E* ~12 kPa.^[33]^ Gilbert *et. al.* went a step further and showed that muscle stem cells propagated in physiologically-relevant, muscle-like rigidities (12 kPa) did not only maintained self-renewal capacity *in vitro*, but also exhibited improved muscle engraftment and functional regeneration in mice *in vivo*.^[34]^ Along similar lines, oligodendrocyte neuro-progenitor cells lose their normal function as their micro-niche stiffens with aging, yet they can functionally rejuvenated when transplanted to a compliant synthetic niche with stiffnesses that mimic the elasticity of young brains.^[35]^ Similarly, compliant healthy lung-like tissue stiffness completely suppresses collagen type I secretion and proliferation of pro-fibrotic, activated fibroblasts.^[5]^ Collectively, these intriguing observations indicate that some cells are inherently sensitive to matrix stiffness. Further, it implies that there might be a unifying biological mechanism(s) that facilitates cells sensing of optimal tissue-specific stiffness cues, possibly through (epi)-genetically imprinted “homeostatic set-points”.^[4,36]^

Our transcriptomic data hints that such regulatory (epi-)genetic mechanisms might be implicated in defining a tensional “homeostatic set-point” in tendon cells. Soft (2 kPa) and rigid (180 kPa) stiffnesses model states of low or elevated matrix tension, respectively. Pairwise comparison of DEGs revealed that stiffness deviations from tendon-like elasticity of *E* ~35 kPa correlated with expression of histone-related genes (H2A, H2B, H3, and H4 family members) **(Figure 3B)**. Although such differential expression may reflect differences in replicative cell-cycle progression, several lines of evidence argue against that.

First, bioinformatics analysis showed that mechanically regulated pathways, i.e. histone modifications and Hippo pathway, were the top two most enriched pathways in the DEGs of intermediate stiffness (35 kPa) compared to the stiffer substrates (180 kPa). Second, 70% of the top over-represented GO terms are related to epigenetic processes that are reported to be involved in stiffness-sensing and mechanical memory **(Figure 3D and G)**. Biophysical cues, including stiffness, can regulate the cells’ “epigenetic states” by altering chromatin organization and accessibility.^[37,38]^ Human mesenchymal stromal cells cultured on soft or stiff microenvironments showed differential responses in histone acetylation and chromatin organization that was dependent on the stiffness cues and the length of exposure to stiff hydrogels.^[39]^ Along the same line, matrix stiffening is implicated in the pathological activation of cardiac stromal cells through a process involving histone deacetylases and chromatin remodeling.^[40]^ In the context of tendon, Heo and colleagues have recently demonstrated that substrate stiffness alters chromatin spatial organization (Histone-H2B nanodomains) of tendon-derived stromal cells that is in agreement with these observations.^[41]^ When tendon fibroblasts were conditioned to soft (3 kPa) methacrylated hyaluronic acid hydrogels or ultra-stiff glass surfaces, histone-H2B nanodomains condensed and relocated to the nuclear periphery. This spatial chromatin condensation was reminiscent to what they observed in cells obtained from diseased tendinopathic or aged tendons. Strikingly, tenocytes that were cultured in substrates with tendon-like stiffness (30 kPa) exhibited diffuse chromatin distribution throughout the nucleus akin to spatial patterning observed in healthy young tendons.^[41]^ While our data does not establish direct causal relationship between substrate stiffness and chromatin architecture, upstream TFs analysis predicted enrichment of several transcriptional effectors known to be involved in epigenetic machineries. Notably, SUZ12 is the only inferred TF that is shared across all the stiffness comparisons **(Figure 3 J-L)**. In the most rigid condition (180 kPa), SUZ12 moves up the list to be the most enriched putative TFs while EZH2 appears among the top 10 most enriched TFs. Both SUZ12 and EZH2 form the core subunits of the polycomb repressive complex 2 (PRC2). PRC2 complex binds remodeled heterochromatin and catalyzes the formation of H3K27me3 epigenetic mark, which is widely acknowledged to play a key role by acting as a “mechanical rheostat” in response to dynamic mechanical loading or in instilling epigenetic mechanical memory in mesenchymal stromal cells.^[38,42]^

Another key insight from this work is we documented the extent of the genome-wide phenotypic drift of freshly isolated tendon stromal cells when maintained in conventional tissue culture plastics (TCP). Progressive cell expansion in ultra-stiff TCP is widely acknowledged as a key driver of spontaneous activation of tenocytes towards pro-fibrotic myofibroblasts. Our data here show that standard TCP-culture significantly alters the native phenotype of tendon cells, even without ever expanding these cells in tissue plastic (i.e. initiating and collecting the cells for downstream analysis at passage 0) **(Figure 3, Supplementary figure S1)**. These observations in agreement with recent findings made by our lab and others.^[25,43]^ van Vijven *et. al.* has shown that expansion of mouse tenocytes in 2D TCP substrates induced loss of native phenotype of mouse tendon-derived cells. Consistently, Imada and colleagues reported similar findings and implicated serum-supplementation in culture medium in mediating osteoblastic-like changes in rat Achilles tenocytes. Interestingly, while the tenogenic markers were strongly suppressed in our TCP conditions, the osteogenic markers *ALPL* (Alkaline phosphatase) and *SPP1* (Osteopontin) were significantly upregulated compared to native tendons in our data **(Figure supplementary S1 A and B)**. One intriguing question is whether this differential expression reflects an underlying differentiation of subpopulation of tendon-resident stem cells or a serum-dependent proliferation ALPL^+^ cells.

Furthermore, the detailed transcriptome analysis revealed that tendon cells in stiff TCP cultures exhibited strong transcriptional signatures suggestive of angiogenesis, fibro-inflammatory (EMT, TGF-β1 pathway), and metabolic (glycolysis) activation. Two independent reports have established that TCP *per se* activates quiescent stromal cells into the myofibroblast phenotype, through coordinated short-term signaling cascades and long-term epigenetic mechanisms.^[24,30]^ PI3k/AKT pathway was implicated in mediating the short-term effects of pathologically-stiff matrix, while we recently uncovered that ERK1/2 kinases are central checkpoints relaying short-term signaling responses to mechanical stress in tendons.^[21, 30, 44]^ ERK1/2 were overrepresented across all stiffness comparison in this work, further reinforcing its role in mediated mechano-responses.^[21]^ Additionally, stiff matrix mechanical cues were shown to rewire the metabolic activity of cells *in vitro* towards “mechanoresponsive glycolysis”, through a mechanism involving cytoskeletal tension, actin organization and proteasomal degradation of the metabolic enzyme phosphofructokinase (PFK).^[45]^

Moreover, increased glucose metabolism, through glycolysis, was reported to modulate the transcriptional activity of YAP/TAZ suggesting that mechanoresponsive glycolysis may instill a self-amplifying feed-forward loop between substrate stiffness and mechanosignaling.^[46]^ Interestingly, our PDMS mechano-variant substrates completely abrogated the angiogenesis, fibro-inflammatory and metabolic signatures when compared to TCP transcriptome **(Figure 3, Figure 4, Supplementary figures S7 and S8)**. Collectively, this suggests that PDMS elastomeric gels do only provide an optimal “tenogenic” stiffness but also mitigate the undesirable side effect of conventional tissue culture plastics.

In summary, we describe a novel PDMS-based culture platform that greatly improves scalability, reproducibility, and ease of handling of mechano-variant substrates. We determined that 35 kPa substrates tend to support tendon marker expression while mitigating activation of pathways associated with loss of tensional homeostasis. Finally, we highlight that conventional TCP culture of tendon cells activates proinflammatory, proangiogenic and metabolic pathways that markedly differ from freshly isolated tendon cells and which brings the scientific probity of their use into question.

## 4. Experimental Section/Methods

### Preparation of silicone mechano-variant substrates

Silicone gels were prepared by mixing pure liquid vinyl-terminated polydimethylsiloxane (PDMS) (1,000 cSt - DMS-VM31, Lot. 1K-39727. Gelest), hydride-terminated PDMS (500 cSt – DMS-HM25, Lot. 5L-26253. Gelest), 7-8% (Methylhydrosiloxane)-dimethylsiloxane copolymer, trimethylsiloxy-terminated (AB146377, abcr GmbH) and Platinum-divinyltetramethyldisiloxane catalyst (AB146697, abcr GmbH). A small amount of 0.2% (*v/v*) 10-Undecenoic acid (124672, Sigma-Aldrich) was added to the mix to allow for direct coupling of primary amines. The gels’ Young moduli were tuned by varying the amounts of crosslinker, vinyl-terminated, and hydride-terminated PDMS in the mix, while keeping the ratio of the platinum catalyst constant at 0.015% (*v/v*). To polymerize the gels, the silicone solution was thoroughly mixed, degassed under vacuum for 30 minutes and allowed to set at room temperature (for a total of one hour). Silicone mixture was then poured into desired cell culture vessels and cured at 70°C for another 23 hours. Cured PDMS substrates were sterilized using UV light and 70-80% ethanol under aseptic conditions.

### PDMS functionalization and ECM coating

To allow for cell adhesion, PDMS substrates were functionalized with in-house prepared rat tail collagen type I using the mild Sulfo-NHS plus EDC (carbodiimide) crosslinking procedure (carboxyl-to-amine). Substrates were washed once with sterile MES (2-(N-morpholino)ethanesulfonic acid) buffer (28390, Thermo Scientific™), then activated with Sulfo-NHS/EDC: (3 mg) EDC plus (3mg) NHS in (1 ml) MES buffer for 20-60 mins at RT. Collagen type I was dilute to 50 μg ml^−1^ in ice cold PBS, and used to coat the PDMS surface overnight at 4°C (at a ligand density of 10 μg cm^−2^).

### Isolation of human tendon-derived stromal cells

Human tendons biopsies were obtained from healthy donors undergoing autograft tendon transfer procedures for the surgical repair of anterior cruciate ligaments.^[47]^ Excess samples that would have otherwise been discarded were collected with informed donor consent in compliance with the requirements of the Declaration of Helsinki, the Swiss Federal Human Research Act (HRA), and Zurich Cantonal Ethics Commission (Approval number: 2015-0089, 2020-01119). Immediately after surgical dissection, tendon samples were placed in (DMEM)/F12 medium (D8437, Sigma-Aldrich) in the operating room and were placed in 4°C. In the cell culture lab, samples were thoroughly washed in PBS to remove blood or attached muscle debris. Tendon tissue was cut into small pieces (approx. 1 mm^3^ cubes). Human tendon-derived stroma cells were isolated by collagenase digestion using collagenase, Type I (17018029, Gibco™) at 37 °C for 6-12h digestion.^[48]^ After digestion, freshly isolated single cells were seeded directly onto PDMS substrates without further expansion on tissue culture flaks (i.e. passage 0). During the experiments, cells were maintained in Dulbecco’s Modified Eagle’s Medium with L-glutamine, 15 mM HEPES and sodium bicarbonate (DMEM/F12 - D8437, Sigma-Aldrich), supplemented with 10% heat-inactivated fetal bovine serum (FBS - 10500, Gibco™), 1% (v/v) Penicillin-Streptomycin (P/S, P0781, Sigma-Aldrich), 1% MEM Non-essential Amino Acid Solution (M7145, SAFC Sigma-Aldrich). Cells were maintained in a culture incubator set at 37 °C and 5% CO_2_, and fresh media were replenished every 3-4 days until cells reached 70-80% confluency.

### RNA extraction

#### Cells in PDMS substrates

Cells were washed with PBS and were lysed *in situ* using Qiagen RLT Plus lysis buffer (1053393, Qiagen). Cell lysates were snap-frozen in liquid nitrogen and stored at −80°C until further use. To extract the RNA, samples were thawed on ice, vortexed for 1 min., and homogenization using QIAshredder columns (79654, Qiagen). Total RNA was isolated using RNeasy Plus Micro Kit (74034, Qiagen), including a genomic DNA removal step with Qiagen gDNA Eliminator spin columns and on-column DNase digestion (12185010, Invitrogen™). RNA concentration was measured using Qubit™ RNA HS Assay Kit (Q32852, Invitrogen™).

#### Human tendon tissues

Control tissues were freshly snap frozen in liquid nitrogen immediately after the surgical operation. On the day of RNA extraction, samples were transferred to a supercooled Spex microvial grinding cylinder (6757C3, SPEX™ SamplePrep) containing 100 μl of GENEzol™ reagent (4402-GZR200 Labforce LSBio). Samples were pulverized in a bath of liquid nitrogen using a cryogenic mill (Spex6775 FreezerMill) for 2-4 milling cycles at 15 cps. The resulting pulverized tissue lysate was further solubilized in an additional (1-1.5) ml GENEzol. Insoluble ECM fragments were disrupted by passing the whole lysate approximately 10 times through a 21G syringe needle, followed by centrifugation at 5000 g for 5 min.

To extract the RNA, cleared homogenate was thawed and thoroughly mixed with (200μl) of Chloroform (102445, Merck) at a mixing ratio of 1:5. After centrifugation (12 000 g, 15 min, 4°C), the RNA containing aqueous phase was carefully pipetted, mixed with an equal volume of 70% EtOH. Further RNA clean-up was carried out using Mini Scale Kit for human tissue (Invitrogen, 12183018), including an on-column DNase digestion (12185010, Invitrogen™), according to the manufacturer’s instructions. RNA amounts were quantified with Qubit™ RNA HS Assay before submitting the samples for RNA-sequencing.

### Genome-wide RNA sequencing (RNA-Seq)

RNA-Seq libraries construction and sequencing were carried out by GENEWIZ® (Leipzig, Germany). Briefly, mRNA integrity and yield were assessed with Agilent Fragment Analyzer (Agilent Technologies, USA). For mechano-variant culture samples, all the samples passed the quality control criteria for sequencing with an RNA quality number (RQN) ≥ 9. RNA-Seq libraries was prepared with poly-A selection enrichment using NEBNext Ultra II Directional RNA Library Prep Kit for Illumina, following manufacturer’s instructions (NEB, Ipswich, MA, USA). For native human tendons, libraries were prepared using a ribosomal RNA depletion protocol using NEBNext rRNA Depletion Kit (Human/Mouse/Rat) and NEBNext Ultra II Directional RNA Library Prep Kit for Illumina (NEB, Ipswich, MA, USA). Libraries were sequenced on the Illumina NovaSeq 6000 instrument in Pair-End (PE) configuration to a sequencing depth of at least 20 million reads.

#### Data preprocessing and bioinformatics analysis

Primary level bioinformatics analysis was carried out using the R package ezRun, which is implemented within the SUSHI freamwork of the Functional Genomics Center Zurich (ETH Zurich and the University of Zurich).^[49]^ Quality of the raw RNAseq data was verified with FastQC and MultiQC (version 1.9).^[50]^ Raw.fastq files were mapped to the human genome reference (Ensembl GRCh38.p10 - Release_91-2018-02-26) with STAR aligner software.^[51]^ Reads counts per gene were calculated using featureCounts function against the reference genome (Ensembl version 78).^[52]^ Differential expression analysis was performed by using the *DESeq2* R package (version 4.0.3) using default parameters.^[53]^ Pathway enrichment analysis of differentially expressed genes or selected subclusters of genes of interest was carried out using *Enrichr* tool.^[16,17]^ Gene lists were interrogated against KEGG, BioPlanet, Reactome and/or WikiPathways databases.

Gene set functional enrichment was performed using two complimentary methods: 1) pre-ranked Gene Set Enrichment Analysis using *fgsea* [v1.16.0]^[54]^, 2) gene set overrepresentation analysis (ORA) implemented within clusterProfiler package [v 3.18.0].^[26]^ For ORA analysis, *p*-values were adjusted for multiple hypothesis testing using hypergeometric overrepresentation testing. Analyses were performed on both up-regulated and down-regulated genes, unless specified other in the figure legends. For tertiary functional analysis of expression signatures, raw read counts were imported into Omics Playground Suite (BigOmics Analytics, Switzerland) implemented in RStudio (v. 1.3.1093).^[55]^ Functional annotations of heatmap clusters were defined based on geneset-level, Fisher-weighted correlation with the reference database Molecular Signatures Database (Hallmark collection, MSigDB).^[27,56]^ Volcano plots were visualized using VolcaNoseR package.^[57]^

#### Transcription factors and kinase enrichment analyses

Upstream regulators (kinases and transcription factors) that are likely responsible for the changes in gene expression were computationally deduced using *Expression2Kinases* toolkit (MacOS, Version 1.6.1207).^[16]^ Top DEGs (up to 3000 genes) were fed into *Expression2Kinases* pipeline which applies enrichment algorithm to predict and rank potential transcription factors regulating the quired DEGs. Next, it construct a protein-protein transcriptional regulatory subnetwork which are seeded into the Kinase Enrichment Analysis (KEA) module.^[58]^ Predicted kinases were mapped to kinome tree dendrogram using Coral tool and phylogenetic information described elsewhere.^[59]^

## Acknowledgments

We would like to thank the donors who generously consented to donate their tendon tissue for research. We also gratefully acknowledge the contribution of clinical teams and research nurses, the administrative staff (particularly Ms. Helen Strebel) and cleaning staff at Balgrist University Hospital, and the Swiss Center for Musculoskeletal Biobanking. We would like to thank Lennart Opitz (Functional Genomics Center, Zürich) for his support on RNA‐ sequencing data. This work has been funded by the Cariplo Foundation [2016-0481], the Vontobel Foundation, and institutional funding of both the ETH Zurich and the University Hospital Balgrist.

## Author contributions

A.A.H and J.G.S conceived the study and designed the experiments. A.A.H, B.N, M.B, and N.G developed experimental protocols and conducted experiments. A.A.H analyzed and interpreted the data and wrote the manuscript. J.G.S acquired funding, supervised the project, interpreted the data, and revised the manuscript. All authors approved the manuscript.

Received: ((will be filled in by the editorial staff))

Revised: ((will be filled in by the editorial staff))

Published online: ((will be filled in by the editorial staff))

## Supplementary Information

**Supplementary figure S1.**
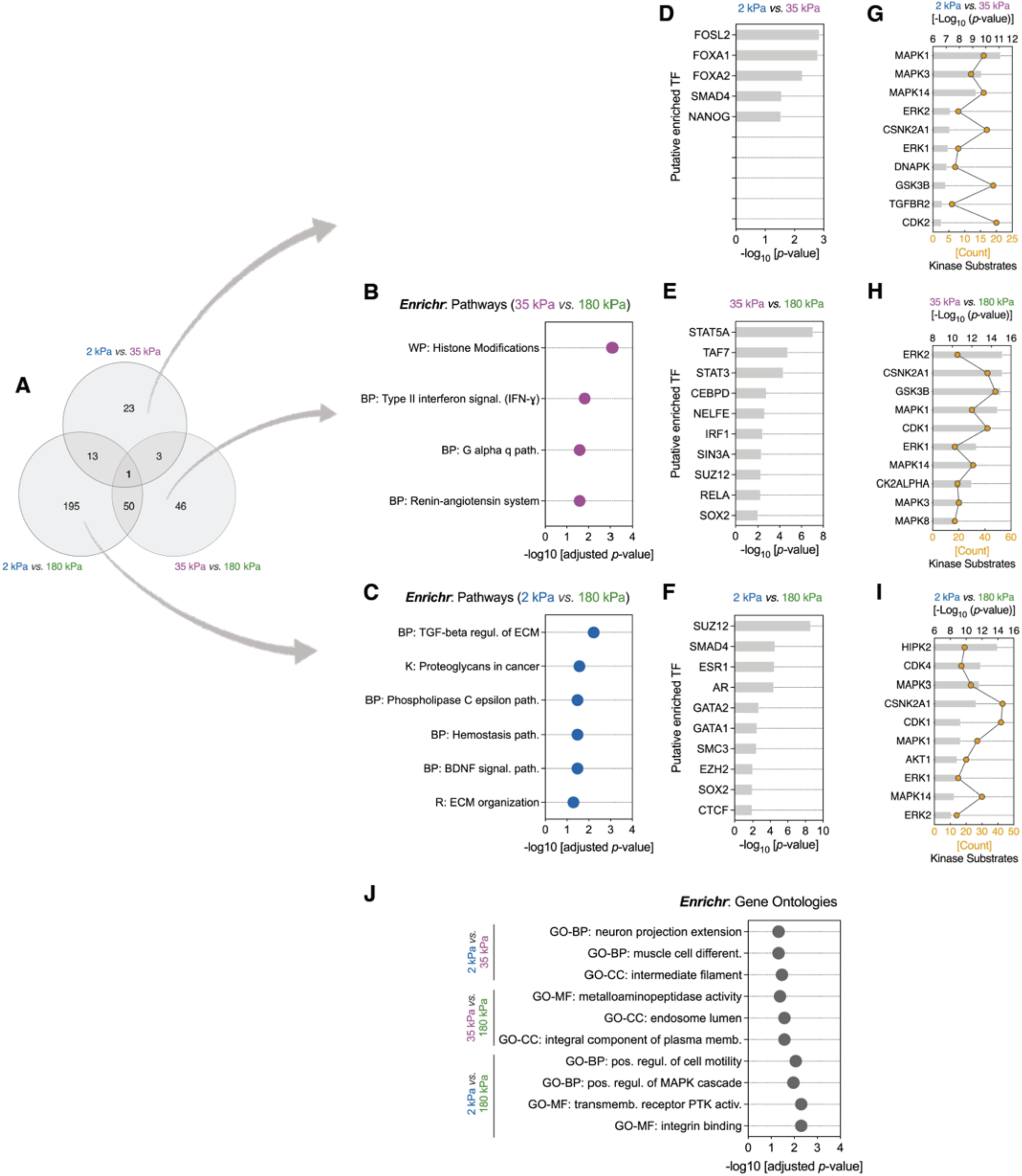
Stiffness-specific enriched pathways and inferred upstream kinases and transcription factors. **(A)** Venn diagram depicting the overlapped significantly expressed genes (DEGs) between different stiffness comparisons. Unique genes to each stiffness comparison (*i.e.* stiffness-specific genes) were used for downstream analysis. **(B-C)** Top enriched pathways for each stiffness-specific comparison; (H) 35kPa *vs.* 180kPa, (E) 2kPa *vs.* 180kPa. Enrichment analysis was performed in *Enrichr.* **(D-F)** Bar plots show the predicted top 5-10 most significantly enriched transcription factors upstream of the stiffness-specific DEGs. Predicted TFs are sorted by significance level (adjusted *p*-value < 0.05). **(G-I)** Bar plots indicate the predicted top 10 most significantly enriched kinases upstream of the stiffness-specific DEGs. Predicted kinases are ranked by significance level (adjusted *p* value < 0.05). TF and kinases were predicted using the TF and Kinase Enrichment Analysis modules of the *Expression2Kinase* tool. **(J)** Stiffness-specific top enriched GO terms. Enrichment analysis was performed in *Enrichr*. Significance achieved at *q-*value < 0.05.

**Supplementary figure S2.**
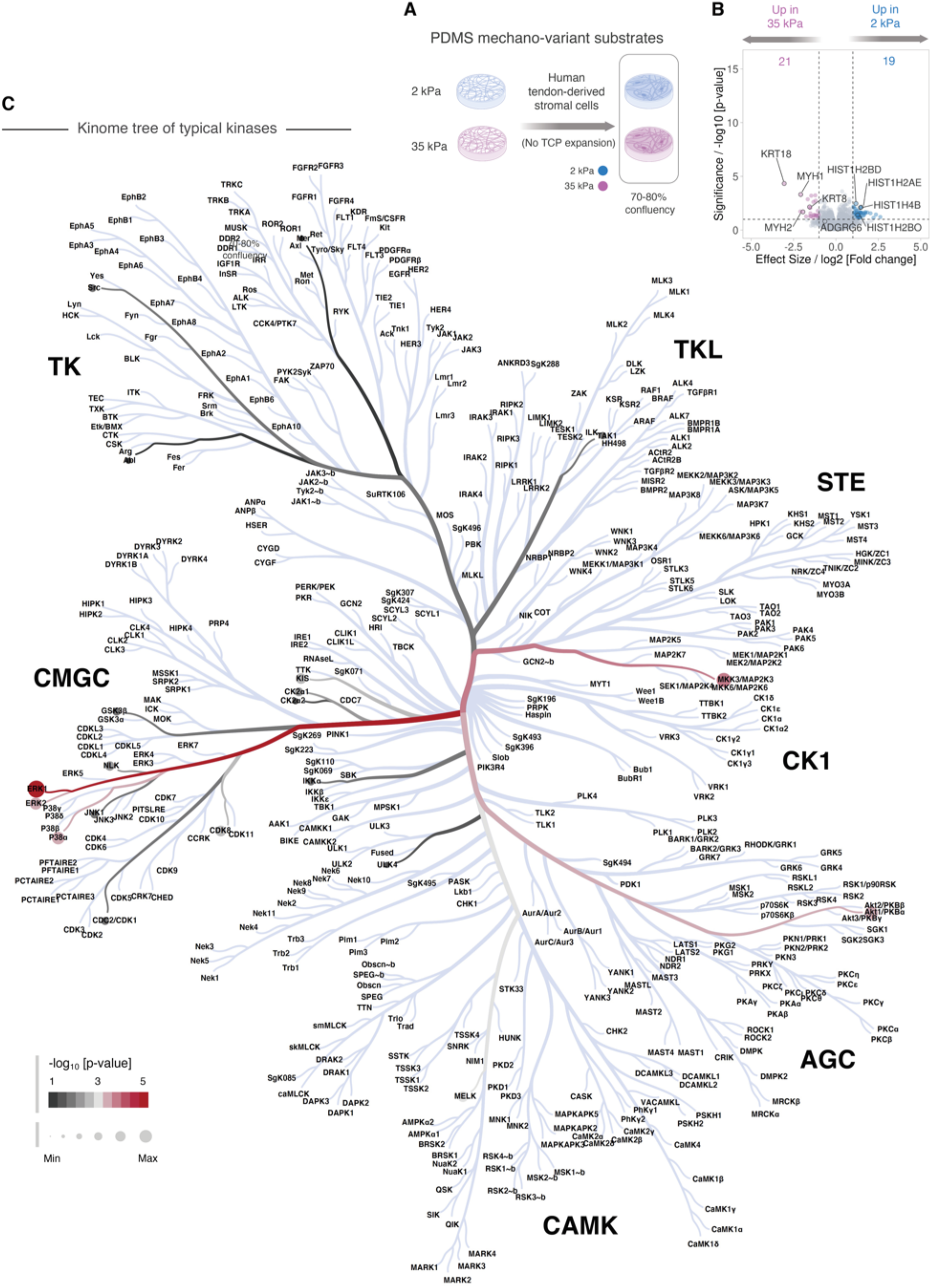
Predicted key signaling kinases upstream of the DEGs in 2 kPa *vs.* 35 kPa PDMS. **(A**) Schematic of the experimental conditions. **(B)** RNA-seq volcano plot of DEGs. Horizontal line corresponding to *p* value ≤0.01 and vertical lines are at a cutoff of log2 [Fold change] ± 1. The same panel also appears in Figure 3 B. **(B)** Kinome tree dendrogram mapping of all the enriched upstream protein kinases in 2 kPa *vs.* 35 kPa PDMS. Circles’ color and size encode the enrichment significance value, with red color reflecting higher significance. Blue branches depict “no value” background.

**Supplementary figure S3.**
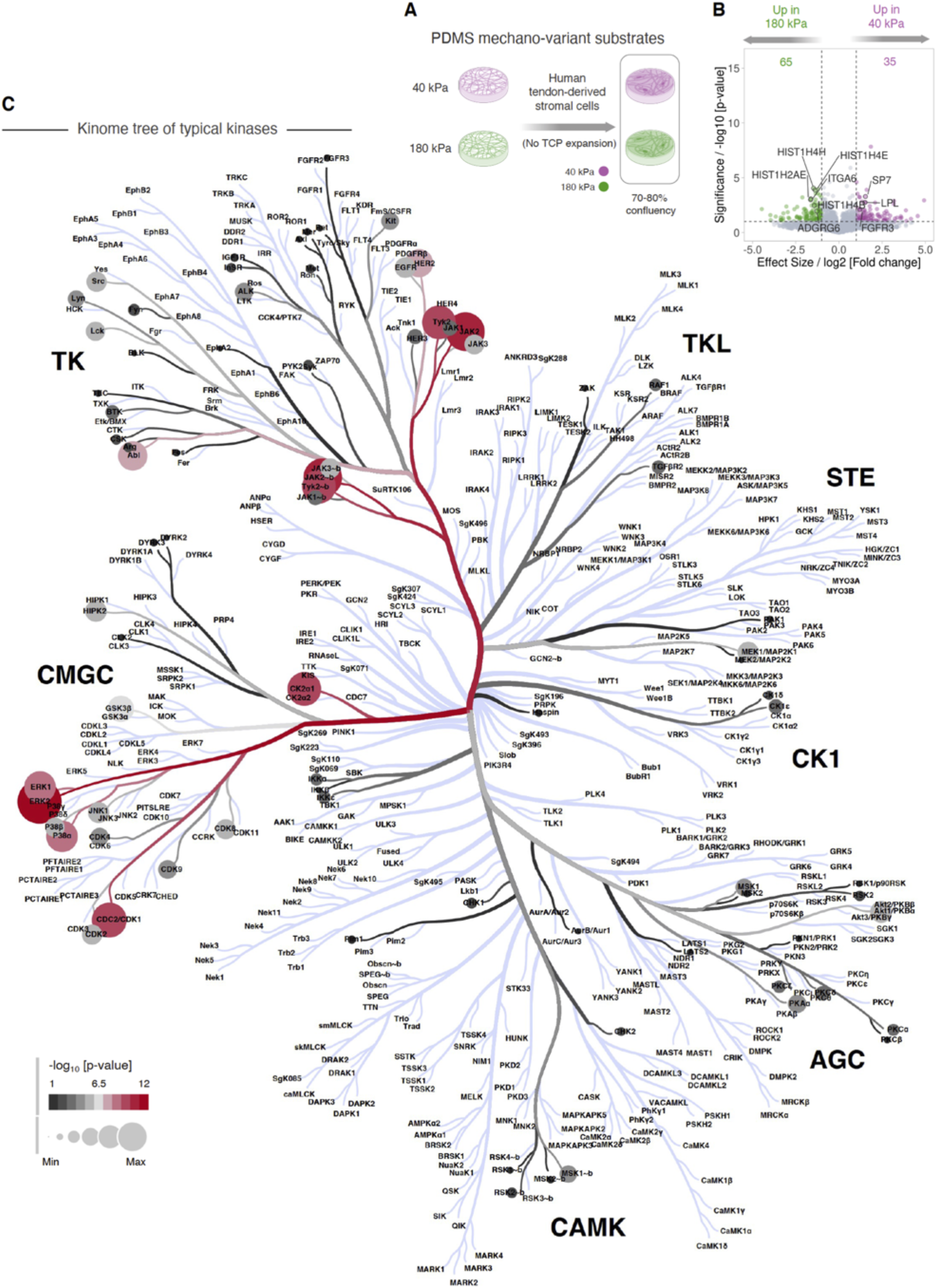
Predicted key signaling kinases upstream of the DEGs in 35 kPa *vs.* 180 kPa PDMS. **(A**) Schematic of the experimental conditions. **(B)** RNA-seq volcano plot of DEGs. Horizontal line corresponding to *p* value ≤0.01 and vertical lines are at a cutoff of log2 [Fold change] ± 1. The same panel also appears in Figure 3 B. **(B)** Kinome tree dendrogram mapping of all the enriched upstream protein kinases in 35 kPa *vs.* 180 kPa PDMS. Circles’ color and size encode the enrichment significance value, with red color reflecting higher significance. Blue branches depict “no value” background.

**Supplementary figure S4.**
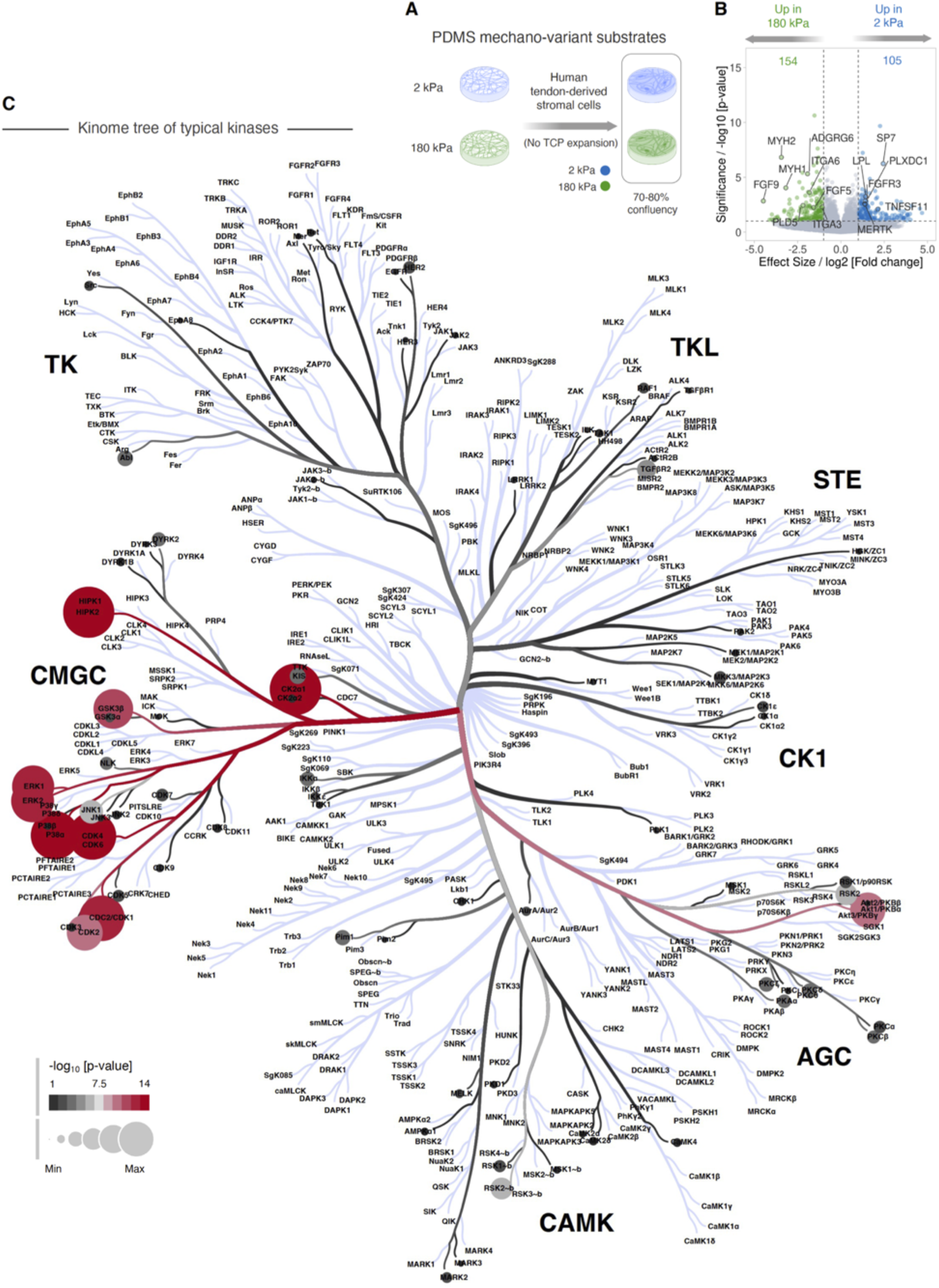
Predicted key signaling kinases upstream of the DEGs in 2 kPa *vs.* 180 kPa PDMS. (**A**) Schematic of the experimental conditions. **(B)** RNA-seq volcano plot of DEGs. Horizontal line corresponding to *p* value ≤0.01 and vertical lines are at a cutoff of log2 [Fold change] ± 1. The same panel also appears in Figure 3 B. **(B)** Kinome tree dendrogram mapping of all the enriched upstream protein kinases in 2 kPa *vs.* 180 kPa PDMS. Circles’ color and size encode the enrichment significance value, with red color reflecting higher significance. Blue branches depict “no value” background.

**Supplementary figure S5.**
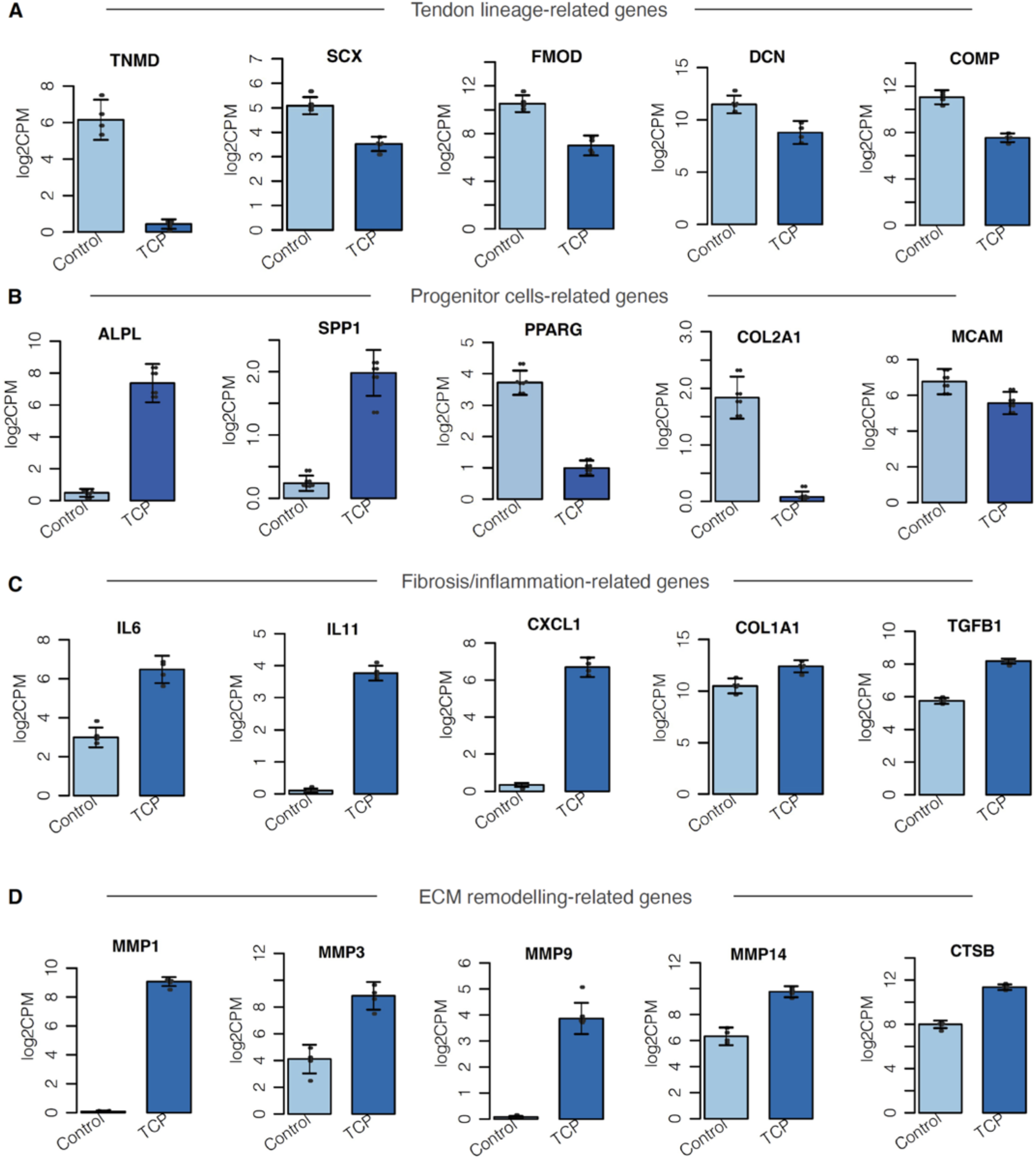
Phenotypic drift is evident in TCP conditioned tendon fibroblasts. RNA-Seq expression values of **(A**) Tendon lineage-related genes, **(B)** Progenitor cells differentiation markers, **(C)** Fibrosis/inflammation-related genes, and **(D)** ECM remodeling-related genes. Control represents native tendon tissue.

**Supplementary figure S6.**
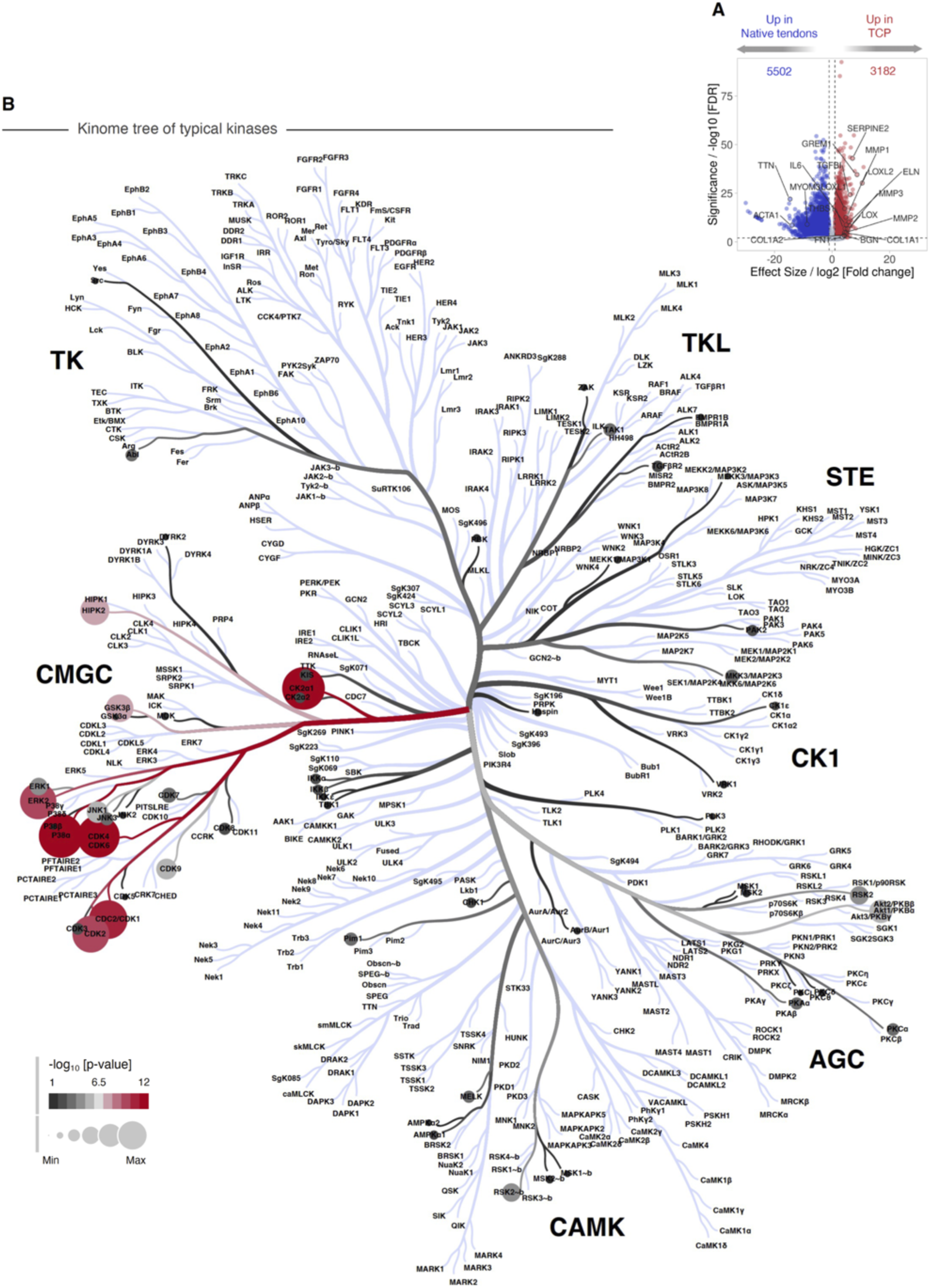
Predicted key signaling kinases upstream of the DEGs in TCP *vs.* Native tendons. **(A**) RNA-seq volcano plot of DEGs. Horizontal line corresponding to an FDR ≤0.01 and vertical lines are at a cutoff of log2[Fold change] ± 1. The same panel also appears in Figure 2C. **(B)** Kinome tree dendrogram mapping of all the enriched upstream protein kinases in TCP *vs.* Native tendons. Circles’ color and size encode the enrichment significance value, with red color reflecting higher significance. Blue branches depict “no value” background.

**Supplementary figure S7.**
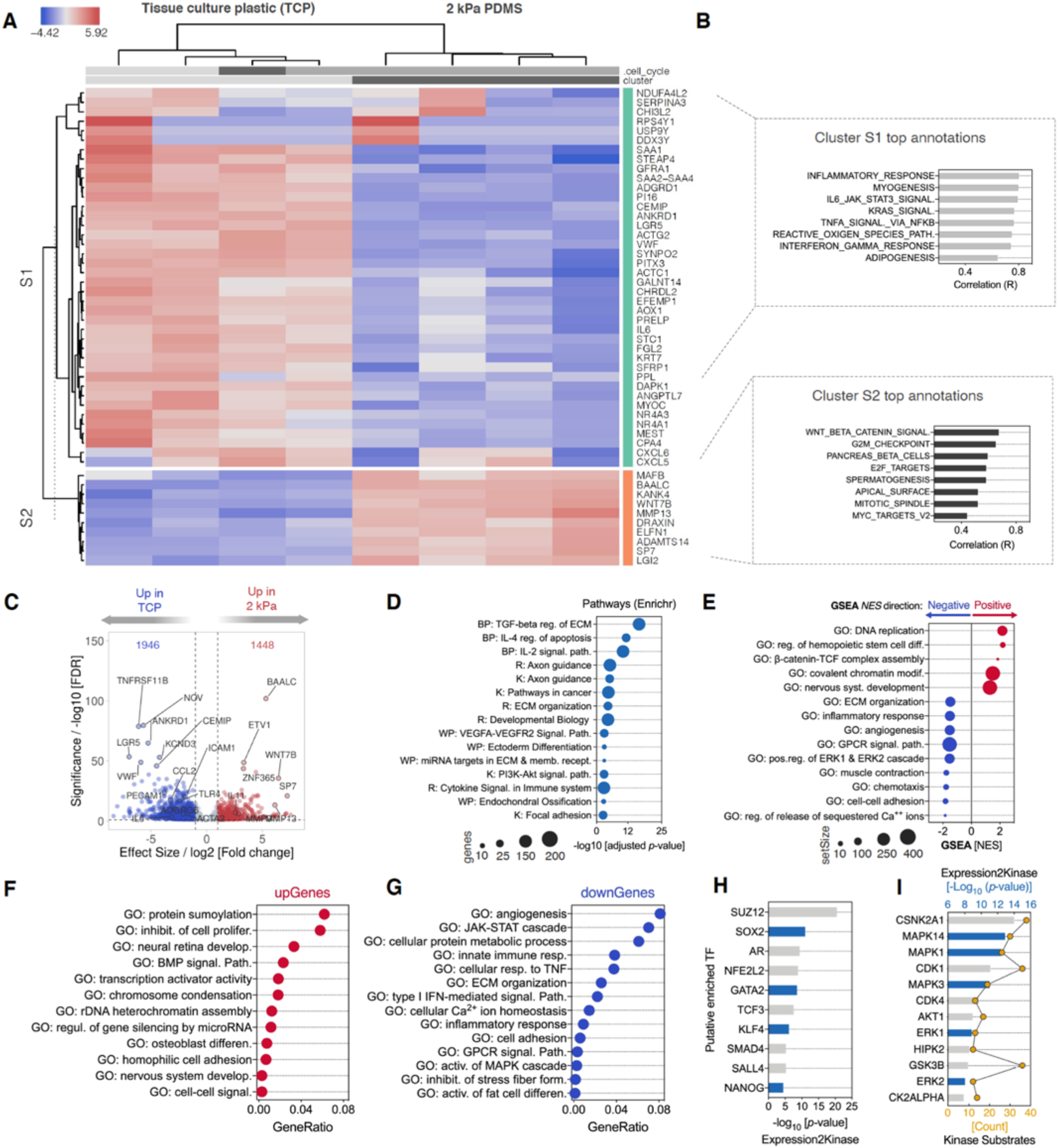
Transcriptome analysis of tendon-derived stromal cells on 2 kPa soft substrates *vs.* TCP. **(A)** Heatmap of gene-level hierarchical clustering of the top 50 differentially expressed genes (DEG) in tendon stromal fibroblasts cultured on soft PDMS substrates (*E.* 2 kPa) *vs.* tissue culture plastic (TCP). Columns represent individual samples (N= 4 biological replicates from different donors). Blue denotes downregulated genes; red denotes upregulated genes. **(B)** Functional annotation of S1 and S2 gene clusters in the heatmap. Bar plots depict the fisher-weighted, average correlations of the cluster with the annotation terms queried against MSigDB Hallmark database. **(C)** RNA-seq volcano plot of DEGs of 2 kPa conditioned human tenocytes relative to TCP control. Colored dots show the 3,394 significantly expressed genes, as determined by *DESeq2* methods, with the horizontal line corresponding to an FDR ≤0.05 and vertical lines are at a cutoff of log2[Fold change] ± 1. **(D)** Enriched pathways analysis of a subset of DEGs using *Enrichr* queried against BioPlanet 2019, Reactome and WikiPathways 2019 Human databases. All hits had an adjusted *p*-value < 0.05. **(E)** Pre-ranked Gene Set Enrichment Analysis (*GSEA*) of positively and negatively enriched biological processes by Normalized Enrichment Score (NES) in 2 kPa cultured cells (Adjusted *p*-value < 0.05). **(F-G)** Overrepresentation analysis (ORA) of Biological Processes GO terms of upregulated (F) and downregulated (G) DEGs. Analysis was performed using a hypergeometric over-representation test against the GO database, with significance cutoff at *q-*value < 0.05. **(H)** Bar plot depicts the predicted top 10 most significantly enriched transcription factors upstream of the DEGs. Predicted TFs are sorted by significance level (adjusted *p*-value < 0.05). **(I)** Top 10 kinases upstream of predicted TFs in (F) were identified using the Kinase Enrichment Analysis module of the *Expression2Kinase* pipeline.

**Supplementary figure S8.**
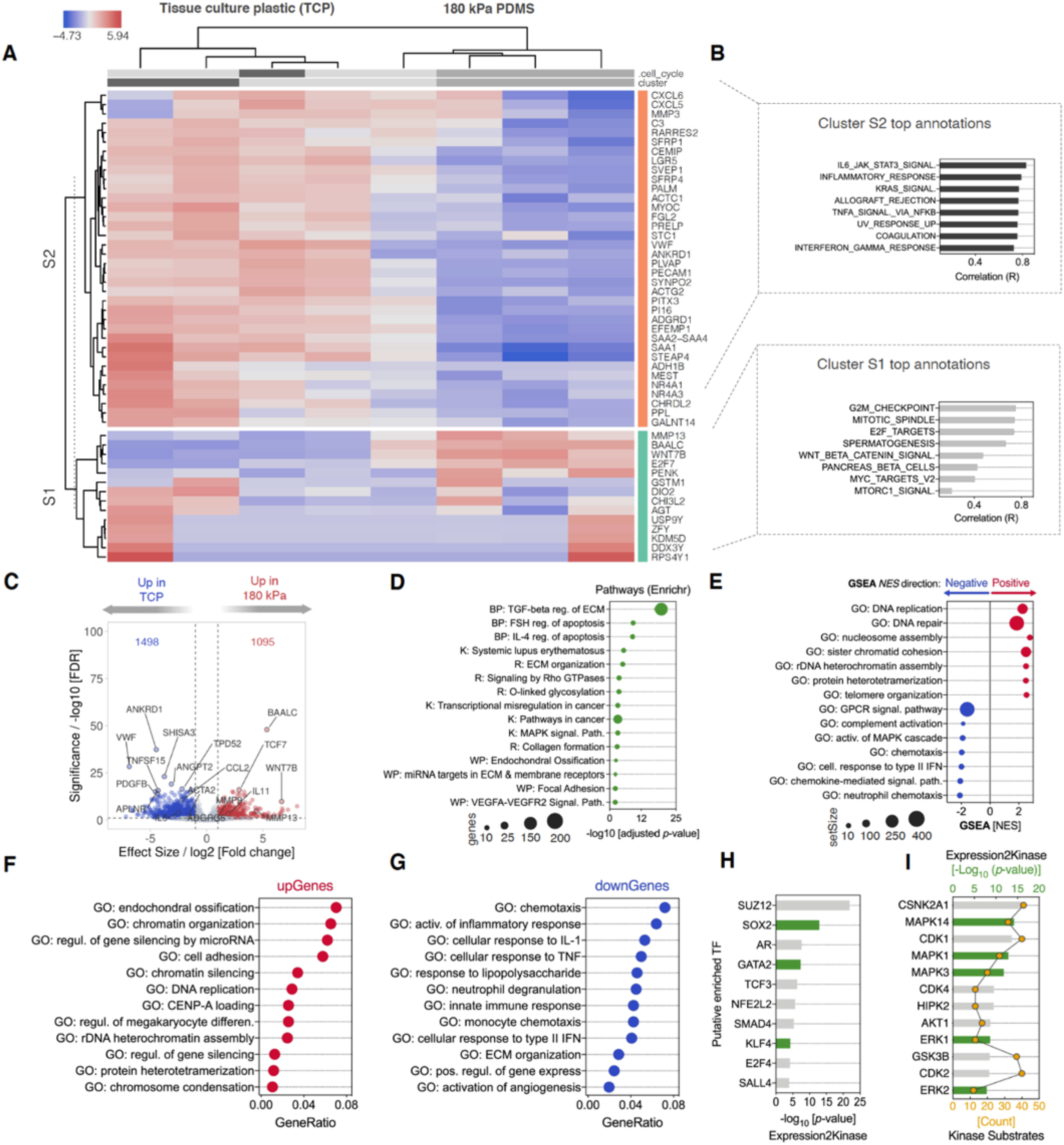
Transcriptome analysis of tendon-derived stromal cells on 180 kPa stiff substrates *vs.* TCP. **(A)** Heatmap of gene-level hierarchical clustering of the top 50 differentially expressed genes (DEG) in tendon stromal fibroblasts cultured on stiff PDMS substrates (*E.* 180 kPa) *vs.* tissue culture plastic (TCP). Columns represent individual samples (N= 4 biological replicates from different donors). Blue denotes downregulated genes; red denotes upregulated genes. **(B)** Functional annotation of S1 and S2 gene clusters in the heatmap. Bar plots depict the fisher-weighted, average correlations of the cluster with the annotation terms queried against MSigDB Hallmark database. **(C)** RNA-seq volcano plot of DEGs of 180 kPa conditioned human tenocytes relative to TCP control. Colored dots show the 2,593 significantly expressed genes, as determined by *DESeq2* methods, with the horizontal line corresponding to an FDR ≤0.05 and vertical lines are at a cutoff of log2[Fold change] ± 1. **(D)** Enriched pathways analysis of a subset of DEGs using *Enrichr* queried against BioPlanet 2019, Reactome and WikiPathways 2019 Human databases. All hits had an adjusted *p*-value < 0.05. **(E)** Pre-ranked Gene Set Enrichment Analysis (*GSEA*) of positively and negatively enriched biological processes by Normalized Enrichment Score (NES) in 180 kPa cultured cells (Adjusted *p*-value < 0.05). **(F-G)** Overrepresentation analysis (ORA) of Biological Processes GO terms of upregulated (F) and downregulated (G) DEGs. Analysis was performed using a hypergeometric over-representation test against the GO database, with significance cutoff at *q-*value < 0.05. **(H)** Bar plot depicts the predicted top 10 most significantly enriched transcription factors upstream of the DEGs. Predicted TFs are sorted by significance level (adjusted *p*-value < 0.05). **(I)** Top 10 kinases upstream of predicted TFs in (F) were identified using the Kinase Enrichment Analysis module of the *Expression2Kinase* pipeline.

**Supplementary Table S1.** Patient donor details and demographics Healthy tissues were obtained from healthy donors undergoing autograft tendon transfer procedures for the surgical repair of anterior cruciate ligaments (ACL).

| Tissue banking ID | Age | Sex | Tendon | Clinical info |
| --- | --- | --- | --- | --- |
| 326 | 27 | F | Semitendinosus | ACL, Non-diabetic |
| 936 | 27 | M | Semitendinosus | ACL, Non-diabetic |
| 964 | 27 | M | Semitendinosus | ACL, Non-diabetic |
| 1002 | 21 | F | Semitendinosus | ACL, Non-diabetic |
| 1010 | 30 | M | Semitendinosus | ACL, Non-diabetic |
| 1024 | 18 | F | Semitendinosus | ACL, Non-diabetic |

## Notes

### Competing Interest Statement

The authors have declared no competing interest.

## References

[1] J. Swift, I. L. Ivanovska, A. Buxboim, T. Harada, P. C. Dingal, J. Pinter, J. D. Pajerowski, K. R. Spinler, J. W. Shin, M. Tewari, F. Rehfeldt, D. W. Speicher, D. E. Discher, Science 2013, 341, 1240104.

[2] C. T. Thorpe, C. P. Udeze, H. L. Birch, P. D. Clegg, H. R. Screen, Eur Cell Mater 2013, 25, 48.

[3] N. Wang, J. D. Tytell, D. E. Ingber, Nature Reviews Molecular Cell Biology 2009, 10, 75.

[4] J. D. Humphrey, E. R. Dufresne, M. A. Schwartz, Nat Rev Mol Cell Biol 2014, 15, 802.

[5] F. Liu, J. D. Mih, B. S. Shea, A. T. Kho, A. S. Sharif, A. M. Tager, D. J. Tschumperlin, Journal of Cell Biology 2010, 190, 693.

[6] M. W. Parker, D. Rossi, M. Peterson, K. Smith, K. Sikstrom, E. S. White, J. E. Connett, C. A. Henke, O. Larsson, P. B. Bitterman, J Clin Invest 2014, 124, 1622.

[7] H. Rubin, Public Health Rep 1966, 81, 843.

[8] R. G. Wells, D. E. Discher, Sci Signal 2008, 1, pe13; C. F. Guimaraes, L. Gasperini, A. P. Marques, R. L. Reis, Nature Reviews Materials 2020, 5, 351.

[9] W. J. Polacheck, C. S. Chen, Nat Methods 2016, 13, 415; W. Xi, T. B. Saw, D. Delacour, C. T. Lim, B. Ladoux, Nature Reviews Materials 2019, 4, 23.

[10] B. Trappmann, J. E. Gautrot, J. T. Connelly, D. G. Strange, Y. Li, M. L. Oyen, M. A. Cohen Stuart, H. Boehm, B. Li, V. Vogel, J. P. Spatz, F. M. Watt, W. T. Huck, Nat Mater 2012, 11, 642; J. H. Wen, L. G. Vincent, A. Fuhrmann, Y. S. Choi, K. C. Hribar, H. Taylor-Weiner, S. Chen, A. J. Engler, Nat Mater 2014, 13, 979.

[11] J. N. Lee, X. Jiang, D. Ryan, G. M. Whitesides, Langmuir 2004, 20, 11684.

[12] G. Vertelov, E. Gutierrez, S. A. Lee, E. Ronan, A. Groisman, E. Tkachenko, Scientific Reports 2016, 6; M. D. A. Norman, S. A. Ferreira, G. M. Jowett, L. Bozec, E. Gentleman, Nature Protocols 2021.

[13] R. I. Sharma, J. G. Snedeker, Biomaterials 2010, 31, 7695.

[14] M. S. Rehmann, J. I. Luna, E. Maverakis, A. M. Kloxin, J Biomed Mater Res A 2016, 104, 1162.

[15] R. I. Sharma, J. G. Snedeker, PLoS One 2012, 7, e31504.

[16] E. Y. Chen, C. M. Tan, Y. Kou, Q. N. Duan, Z. C. Wang, G. V. Meirelles, N. R. Clark, A. Ma’ayan, Bmc Bioinformatics 2013, 14.

[17] M. V. Kuleshov, M. R. Jones, A. D. Rouillard, N. F. Fernandez, Q. N. Duan, Z. C. Wang, S. Koplev, S. L. Jenkins, K. M. Jagodnik, A. Lachmann, M. G. McDermott, C. D. Monteiro, G. W. Gundersen, A. Ma’ayan, Nucleic Acids Research 2016, 44, W90.

[18] Z. Xie, A. Bailey, M. V. Kuleshov, D. J. B. Clarke, J. E. Evangelista, S. L. Jenkins, A. Lachmann, M. L. Wojciechowicz, E. Kropiwnicki, K. M. Jagodnik, M. Jeon, A. Ma’ayan, Current Protocols 2021, 1, e90.

[19] E. Y. Chen, H. Xu, S. Gordonov, M. P. Lim, M. H. Perkins, A. Ma’ayan, Bioinformatics 2012, 28, 105.

[20] G. V. Shivashankar, Annual Review of Biophysics, Vol 40 2011, 40, 361; K. A. Jansen, D. M. Donato, H. E. Balcioglu, T. Schmidt, E. H. J. Danen, G. H. Koenderink, Biochimica Et Biophysica Acta-Molecular Cell Research 2015, 1853, 3043; L. Irons, J. D. Humphrey, Plos Computational Biology 2020, 16.

[21] U. Blache, S. L. Wunderli, A. A. Hussien, T. Stauber, G. Flückiger, M. Bollhalder, B. Niederöst, S. F. Fucentese, J. G. Snedeker, Scientific Reports 2021, 11, 6838.

[22] J. L. Balestrini, S. Chaudhry, V. Sarrazy, A. Koehler, B. Hinz, Integrative Biology 2012, 4, 410.

[23] C. Yang, M. W. Tibbitt, L. Basta, K. S. Anseth, Nat Mater 2014, 13, 645.

[24] C. X. Li, N. P. Talele, S. Boo, A. Koehler, E. Knee-Walden, J. L. Balestrini, P. Speight, A. Kapus, B. Hinz, Nat Mater 2017, 16, 379.

[25] L. Yao, C. S. Bestwick, L. A. Bestwick, N. Maffulli, R. M. Aspden, Tissue Eng 2006, 12, 1843.

[26] G. C. Yu, L. G. Wang, Y. Y. Han, Q. Y. He, Omics-a Journal of Integrative Biology 2012, 16, 284.

[27] A. Subramanian, P. Tamayo, V. K. Mootha, S. Mukherjee, B. L. Ebert, M. A. Gillette, A. Paulovich, S. L. Pomeroy, T. R. Golub, E. S. Lander, J. P. Mesirov, Proceedings of the National Academy of Sciences of the United States of America 2005, 102, 15545.

[28] J. Reimand, R. Isserlin, V. Voisin, M. Kucera, C. Tannus-Lopes, A. Rostamianfar, L. Wadi, M. Meyer, J. Wong, C. Xu, D. Merico, G. D. Bader, Nat Protoc 2019, 14, 482.

[29] K. A. Rosowski, E. Vidal-Henriquez, D. Zwicker, R. W. Style, E. R. Dufresne, Soft Matter 2020, 16, 5892.

[30] H. Wang, M. W. Tibbitt, S. J. Langer, L. A. Leinwand, K. S. Anseth, Proc Natl Acad Sci U S A 2013, 110, 19336.

[31] B. K. Connizzo, A. J. Grodzinsky, Journal of Biomechanics 2017, 54, 11; B. R. Freedman, A. B. Rodriguez, C. D. Hillin, S. N. Weiss, B. Han, L. Han, L. J. Soslowsky, Journal of the Royal Society Interface 2018, 15; M. Kammoun, R. Ternifi, V. Dupres, P. Pouletaut, S. Meme, W. Meme, F. Szeremeta, J. Landoulsi, J. M. Constans, F. Lafont, M. Subramaniam, J. R. Hawse, S. F. Bensamoun, Scientific Reports 2019, 9; B. M. Chen, X. Cheng, E. W. Dorthe, Y. H. Zhao, D. D’Lima, G. M. Bydder, S. R. Liu, J. Du, Y. J. Ma, Nmr in Biomedicine 2019, 32; H. Xu, T. Liang, L. Wei, J. C. Zhu, X. Liu, C. C. Ji, B. Liu, Z. P. Luo, J Biomech 2021, 116, 110248.

[32] D. E. Discher, P. Janmey, Y. L. Wang, Science 2005, 310, 1139.

[33] A. J. Engler, M. A. Griffin, S. Sen, C. G. Bonnemann, H. L. Sweeney, D. E. Discher, J Cell Biol 2004, 166, 877.

[34] P. M. Gilbert, K. L. Havenstrite, K. E. G. Magnusson, A. Sacco, N. A. Leonardi, P. Kraft, N. K. Nguyen, S. Thrun, M. P. Lutolf, H. M. Blau, Science 2010, 329, 1078.

[35] M. Segel, B. Neumann, M. F. E. Hill, I. P. Weber, C. Viscomi, C. Zhao, A. Young, C. C. Agley, A. J. Thompson, G. A. Gonzalez, A. Sharma, S. Holmqvist, D. H. Rowitch, K. Franze, R. J. M. Franklin, K. J. Chalut, Nature 2019, 573, 130.

[36] K. D. Webster, W. P. Ng, D. A. Fletcher, Biophysical Journal 2014, 107, 146; J. Ye, R. Medzhitov, Nature Metabolism 2019, 1, 947.

[37] T. L. Downing, J. Soto, C. Morez, T. Houssin, A. Fritz, F. L. Yuan, J. L. Chu, S. Patel, D. V. Schaffer, S. Li, Nature Materials 2013, 12, 1154; S. W. Crowder, V. Leonardo, T. Whittaker, P. Papathanasiou, M. M. Stevens, Cell Stem Cell 2016, 18, 39.

[38] S. J. Heo, S. D. Thorpe, T. P. Driscoll, R. L. Duncan, D. A. Lee, R. L. Mauck, Scientific Reports 2015, 5.

[39] A. R. Killaars, J. C. Grim, C. J. Walker, E. A. Hushka, T. E. Brown, K. S. Anseth, Advanced Science 2019, 6.

[40] C. J. Walker, C. Crocini, D. Ramirez, A. R. Killaars, J. C. Grim, B. A. Aguado, K. Clark, M. A. Allen, R. D. Dowell, L. A. Leinwand, K. S. Anseth, Nature Biomedical Engineering 2021.

[41] S.-J. Heo, S. Thakur, X. Chen, C. Loebel, B. Xia, R. McBeath, J. A. Burdick, V. B. Shenoy, R. L. Mauck, M. Lakadamyali, bioRxiv 2021, 2021.04.27.441596.

[42] K. H. Hansen, A. P. Bracken, D. Pasini, N. Dietrich, S. S. Gehani, A. Monrad, J. Rappsilber, M. Lerdrup, K. Helin, Nature Cell Biology 2008, 10, 1291; H. Q. Le, S. Ghatak, C. Y. C. Yeung, F. Tellkamp, C. Gunschmann, C. Dieterich, A. Yeroslaviz, B. Habermand, A. Pombo, C. M. Niessen, S. A. Wickstrom, Nature Cell Biology 2016, 18, 864; R. Leicher, E. J. Ge, X. C. Lin, M. J. Reynolds, W. J. Xie, T. Walz, B. Zhang, T. W. Muir, S. X. Liu, Proceedings of the National Academy of Sciences of the United States of America 2020, 117, 30465.

[43] K. Imada, M. Taniguchi, T. Sato, T. I. Kosaka, K. Yamamoto, A. Ito, Journal of Tokyo Medical University 2013, 71, 143; M. van Vijven, S. L. Wunderli, K. Ito, J. G. Snedeker, J. Foolen, J Orthop Res 2020.

[44] J. L. Young, K. Kretchmer, M. G. Ondeck, A. C. Zambon, A. J. Engler, Scientific Reports 2014, 4; O. Chaudhuri, S. T. Koshy, C. Branco da Cunha, J. W. Shin, C. S. Verbeke, K. H. Allison, D. J. Mooney, Nat Mater 2014, 13, 970; L. W. Liu, Z. F. You, H. S. Yu, L. Zhou, H. Zhao, X. J. Yan, D. L. Li, B. J. Wang, L. Zhu, Y. Z. Xu, T. Xia, Y. Shi, C. Y. Huang, W. Hou, Y. N. Du, Nature Materials 2017, 16, 1252.

[45] J. S. Park, C. J. Burckhardt, R. Lazcano, L. M. Solis, T. Isogai, L. Q. Li, C. S. Chen, B. N. Gao, J. D. Minna, R. Bachoo, R. J. DeBerardinis, G. Danuser, Nature 2020, 578, 621.

[46] E. Enzo, G. Santinon, A. Pocaterra, M. Aragona, S. Bresolin, M. Forcato, D. Grifoni, A. Pession, F. Zanconato, G. Guzzo, S. Bicciato, S. Dupont, Embo Journal 2015, 34, 1349.

[47] R. C. Poulsen, A. J. Carr, P. A. Hulley, Endocrinology 2011, 152, 503.

[48] K. Phelan, K. M. May, Current Protocols in Cell Biology 2015, 66, 1.1.1.

[49] M. Hatakeyama, L. Opitz, G. Russo, W. H. Qi, R. Schlapbach, H. Rehrauer, Bmc Bioinformatics 2016, 17.

[50] P. Ewels, M. Magnusson, S. Lundin, M. Kaller, Bioinformatics 2016, 32, 3047.

[51] A. Dobin, C. A. Davis, F. Schlesinger, J. Drenkow, C. Zaleski, S. Jha, P. Batut, M. Chaisson, T. R. Gingeras, Bioinformatics 2013, 29, 15.

[52] Y. Liao, G. K. Smyth, W. Shi, Bioinformatics 2014, 30, 923.

[53] M. I. Love, W. Huber, S. Anders, Genome Biology 2014, 15.

[54] G. Korotkevich, V. Sukhov, N. Budin, B. Shpak, M. N. Artyomov, A. Sergushichev, bioRxiv 2021, 060012.

[55] M. Akhmedov, A. Martinelli, R. Geiger, I. Kwee, NAR Genomics and Bioinformatics 2019, 2.

[56] A. Liberzon, C. Birger, H. Thorvaldsdottir, M. Ghandi, J. P. Mesirov, P. Tamayo, Cell Systems 2015, 1, 417.

[57] J. Goedhart, M. S. Luijsterburg, Scientific Reports 2020, 10.

[58] A. Lachmann, A. Ma’ayan, Bioinformatics 2009, 25, 684.

[59] G. Manning, D. B. Whyte, R. Martinez, T. Hunter, S. Sudarsanam, Science 2002, 298, 1912; K. S. Metz, E. M. Deoudes, M. E. Berginski, I. Jimenez-Ruiz, B. A. Aksoy, J. Hammerbacher, S. M. Gomez, D. H. Phanstiel, Cell Syst 2018, 7, 347.

